# Recovering biological structure in sparse single-cell proteomics with GIRAFI

**DOI:** 10.64898/2026.05.18.726081

**Authors:** Huan Zhong, Shuxin Chi, Rachel Wong, Jason Rogalski, Ziming Wang, Susanna Chan, Melanie L. Bailey, Arpa Ebrahimi, Gabrielle Jayme, Jerry Yin, Albie Gong, Terrance P. Snutch, Claudia S. Maier, Marco A. Marra, Leonard J. Foster, Xin Tang

**Affiliations:** Department of Biochemistry and Molecular Biology, University of British Columbia, Vancouver, British Columbia V6T 1Z4, Canada; Michael Smith Laboratories, University of British Columbia, Vancouver, British Columbia V6T 1Z4, Canada; BC Cancer Research Institute, BC Cancer, Vancouver, British Columbia V5Z 1L3, Canada; Life Sciences Institute, University of British Columbia, Vancouver, British Columbia V6T 1Z4, Canada; Department of Computer Science, University of British Columbia, Vancouver, British Columbia V6T 1Z4, Canada; Department of Mathematics, University of British Columbia, Vancouver, British Columbia V6T 1Z4, Canada; Department of Chemistry, Oregon State University, Corvallis, Oregon 97331, USA; Department of Cellular and Physiological Sciences, University of British Columbia, Vancouver, British Columbia V6T 1Z3, Canada; Djavad Mowafaghian Centre for Brain Health, University of British Columbia, Vancouver, British Columbia V6T 1Z4, Canada; Linus Pauling Institute, Oregon State University, Corvallis, Oregon 97331, USA; Department of Medical Genetics, University of British Columbia, Vancouver, British Columbia, Canada

## Abstract

Single-cell proteomics (SCP) based on liquid-chromatography mass-spectrometry resolves protein-level cellular heterogeneity, but interpretation remains limited by detection-linked sparsity. SCP profiles continuous, peptide-derived intensities and has lower throughput than single-cell RNA sequencing, making denoising methods for large-scale, count-based transcriptomics difficult to apply. Here we present GIRAFI, a graph-informed statistical learning framework that imputes missing values and reveals reproducible cell states by constraining inference to dataset-aware, prior-knowledge-informed protein neighborhoods. We evaluated GIRAFI across SCP datasets spanning diverse biological/technical contexts. In masking-based recovery experiments and cell-type-specific protein-protein interaction inference, GIRAFI outperformed existing methods, and matched bulk proteomics comparisons corroborated recovery accuracy and ablations supported the graph-informed design. Beyond reduced replicate- and source-associated technical structure, GIRAFI recovered ground-truth cell-type annotations, improved cell state-resolved pathway analysis, and enabled trajectory inference consistent with known time courses. These results establish graph-constrained imputation as an effective strategy for improving SCP robustness, biological structure, interpretation, and cross-dataset comparability.

---

Single-cell proteomics (SCP) brings single-cell analysis closer to phenotype by measuring proteins, the molecules that facilitate signaling, metabolic, and structural programs^1–4^. However, the potential of liquid-chromatography mass-spectrometry (LC-MS)-based SCP is constrained by how data is generated^5–9^. Proteins near the instrument’s detection limit are only observed in a subset of cells^2,10,11^. Coverage also varies with acquisition depth and experimental batches. Beyond these limitations, current workflows still profile far fewer cells than the atlases now common in single-cell RNA-sequencing (scRNA-seq)^12^. SCP results in a small and sparse matrix of continuous peptide intensities^13–15^. Under these conditions, biological readouts, including separation of cell states, marker discovery, pathway activity, and reproducibility across experiments, become unreliable^9,16,17^. The practical question for SCP is therefore whether incomplete measurements can still support reproducible biological inference.

In the early stages of scRNA-seq development, data sparsity was a similar limitation^18–20^, and the field has since produced a rich set of denoising methods to address this issue^21–23^. Denoising SCP is not simply the same problem in another data modality; many scRNA-seq denoising methods were developed for count-based data^24,25^, whereas SCP records continuous peptide- or protein-level intensities. As a result, count-based statistical models such as negative binomial^26^ or zero-inflated negative binomial^27^ models are poorly suited to SCP data. At its core, denoising methods work by borrowing information from recurring structure in the data, often through similarities among cells. In SCP, such cell-cell relationships can be difficult to estimate because datasets are typically smaller, often sampling hundreds rather than tens of thousands of cells in scRNA-seq, and each cell contains many missing values. In this setting, two cells may appear similar only because they share detected proteins from an already limited set. Sparse sampling also makes it difficult for heavily parameterized deep-learning models to learn stable structure from the data alone. Using information across such fragile cell-level neighborhoods can amplify batch effects or overlook rare cell states.

These limitations motivate a denoising strategy that does not rely solely on cell-cell neighborhoods learned from sparse measurements. Proteomics data can be interpreted through decades of curated knowledge on protein interactions, complex assembly, and pathway organization, which together offer a complementary source of data structure^28–30^. Coordinated variation among functionally related proteins, such as proteins within the same pathway or complex, can still be detected across cells even when each individual cell has limited protein coverage. Meanwhile, such priors are not a substitute for the experiment. Global interaction resources are incomplete, denser around well-studied proteins and often silent about which links matter in a particular cell type, perturbation, or time point. In SCP, prior knowledge on protein-protein relationships is therefore most useful when it acts as a light, interpretable constraint, while the measurements themselves determine which relationships are active in the dataset at hand.

We introduce GIRAFI (Graph-Informed Random-Forest Imputation), a statistical learning framework that integrates data-derived and prior-knowledge graphs for denoising sparse SCP data (**Fig. 1a**). GIRAFI leverages protein-protein interactions (PPI) inferred from SCP data together with curated biological knowledge to guide imputation. For each target protein, it learns prediction rules from the observed data while restricting candidate predictors to proteins within defined interaction neighborhoods, thereby ensuring that information is borrowed locally in a biologically relevant manner (see **Methods**). By combining the knowledge priors with data-driven learning, GIRAFI produces imputations that are robust to high levels of sparsity and better aligned with underlying biological organization. This protein-specific design is well-suited to SCP, where informative relationships are often nonlinear, vary from one protein to another, and must be estimated from datasets with limited numbers of cells. GIRAFI achieves state-of-the-art performance across a diverse collection of SCP datasets spanning tissues, conditions, and species. Comprehensive benchmarking shows that GIRAFI improves recovery of masked values, enhances cell-type identification in controlled mixtures, strengthens agreement with matched bulk proteomes, and preserves reproducible biological structure across replicates, data sources, perturbations, and time-course settings. Collectively, these results establish GIRAFI as a practical framework for recovering biologically meaningful structure from sparse SCP data.

**Fig. 1.**
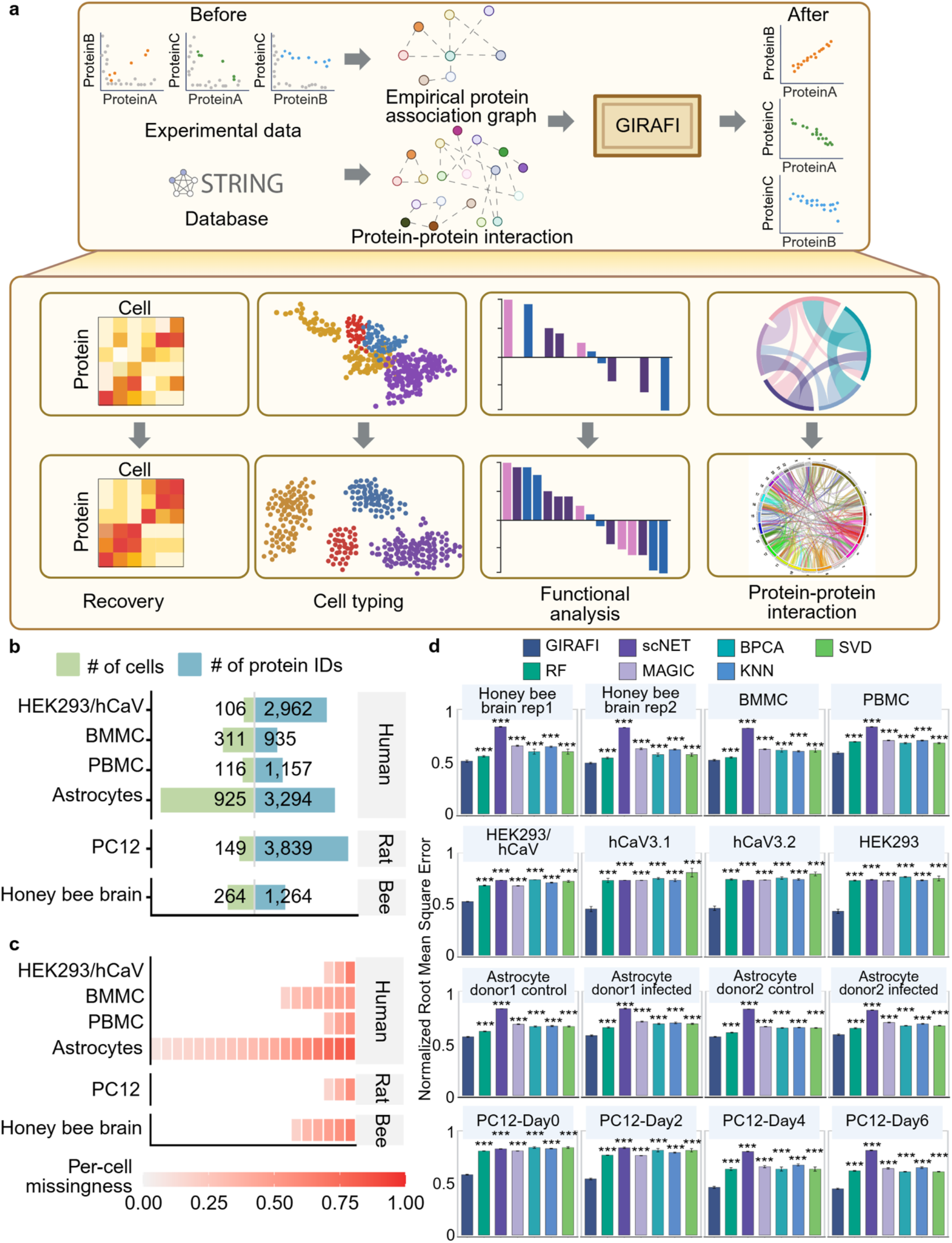
GIRAFI workflow and overview of SCP datasets. **a, Schematic of GIRAFI**. GIRAFI takes a sparse protein-by-cell SCP intensity matrix as input and produces a denoised matrix. The schematic illustrates GIRAFI’s dataset-aware and prior-knowledge-informed strategy. GIRAFI can recover missing values in SCP data, improving downstream cell typing, pathway analyses, and PPI inference. **b, Summary of SCP datasets used for GIRAFI benchmarking**. Bars report the number of cells and quantified proteins for each dataset. **c, Per-cell missingness distributions for each dataset**. After quality control, each dataset is shown as a series of equal-width tiles (each tile represents 25 cells), colored by per-cell missingness (fraction of proteins not detected in a cell; scale shown below). **d, Benchmarking of GIRAFI on a masked values recovery task**. Bar plots summarize mean normalized root mean squared error (NRMSE, see **Methods**) from a standardized mask-and-recover benchmark across the SCP datasets described in **b** and **c**. Within each panel, the x-axis shows the compared recovery methods, encoded by color according to the legend above the plots, and the y-axis shows NRMSE between the held-out masked values and the recovered values. Baseline methods include Random Forest (RF)^33^, scNET^21^, MAGIC (Markov Affinity-based Graph Imputation of Cells)^23^, BPCA (Bayesian Principal Component Analysis)^32^, kNN (k-Nearest Neighbors)^31^, and SVD (Singular Value Decomposition)^31^. Bars show mean ± SD across 5 independent masking replicates; lower NRMSE indicates more accurate recovery. Statistical comparisons were performed using two-sided Welch’s t-tests with GIRAFI as the reference method. Significance is indicated by brackets and asterisks as follows: * p < 0.05, ** p < 0.01, *** p < 0.001.

## Results

### Integrating data-aware and prior-knowledge-informed protein-protein correlation graph for SCP denoising with GIRAFI

Accurate denoising of sparse SCP data requires both data-aware and prior-informed protein structure. GIRAFI models protein relationships with two complementary graphs: an empirical protein association graph estimated from the observed data and a curated PPI prior derived from external knowledge. The empirical graph captures covariance patterns present in the data, whereas the curated PPI constrains inference when sparse detection makes these relationships difficult to estimate robustly from data alone. GIRAFI integrates the two graphs into a fused protein neighborhood graph, in which dataset-specific associations remain dominant while prior knowledge provides biologically plausible regularization. Using this fused graph, GIRAFI denoises each target protein with graph-restricted random forest predictors, either from neighborhood summary statistics or directly from neighboring protein abundances (**Fig. 1a** and **Methods**). This design improves robustness under sparse proteome coverage while preserving experiment-specific structure.

### Systematic evaluation demonstrates state-of-the-art SCP denoising accuracy

We evaluated GIRAFI across diverse SCP datasets spanning multiple species, organs, and experimental designs, including human cell-line mixtures, immune cell datasets, primary astrocyte perturbations, rat PC12 cell-line differentiation time courses, and honey bee brains (**Fig. 1b**). These datasets vary substantially in cell-type composition, proteome coverage, and per-cell missingness (**Fig. 1c**), thereby providing a broad and challenging test bed for SCP denoising method assessment. We compared GIRAFI against representative baselines spanning local-similarity, low-rank, tree-based, diffusion-based, and graph-based imputation strategies, including kNN and SVD^31^, BPCA^32^, Random Forest/missForest^33^, MAGIC^23^, and scNET^21^. Because SCP lacks a complete ground-truth reference at single-cell resolution, a standardized mask-and-recover strategy was used, in which a relatively complete subset of each observed matrix was used to randomly mask a fraction of known values, after which each imputation method was applied and recovered values were compared to the original measurements^34^. Reconstruction error was quantified using Normalized Root Mean Squared Error (NRMSE) and Root Mean Squared Error (RMSE). Across datasets and repeated masking experiments, GIRAFI consistently achieved the lowest NRMSE and RMSE values compared to previous baseline methods (**Fig. 1d** and **Supplementary Fig. 1**). Using mask-and-recover validation, we optimized the graph-fusion parameter λ separately for each astrocyte donor-condition subset. Across the four subsets, the best-performing values were consistently in a low-to-moderate range, indicating that GIRAFI primarily relies on data-derived protein associations while using curated PPI information as a modest prior to stabilize inference in sparse SCP matrices (**Supplementary Fig. 2**).

### GIRAFI removes batch effect across replicates and reveals reproducible cell state structure in *Apis mellifera* forager brain SCP data

We next evaluated GIRAFI in a non-model single-cell proteomics dataset comprising two biological replicates of dissociated *Apis mellifera* forager brains (264 cells and 1,264 quantified proteins after quality control). The data exhibit substantial data sparsity (**Supplementary Fig. 3a**) and replicate-associated batch effects, limiting cross-replicate comparability. Because standardized cell-type annotation is limited in this system, we first performed unsupervised clustering independently within each replicate to obtain replicate-specific cell clusters. We then compared clusters across replicates before and after GIRAFI imputation.

Application of GIRAFI increased effective per-cell protein coverage and reduced cell-to-cell dispersion (**Supplementary Figs. 3 and 4**). Consistent with these improvements, the low-dimensional embedding showed reduced replicate-driven batch effects after imputation (**Fig. 2a**). To quantify this, we computed Spearman correlations between averaged protein profiles for each cluster. In the original data, clusters did not align consistently across replicates. After GIRAFI, clusters exhibited a clear correlation across replicates (**Fig. 2b**). This was also reflected by an increase in the number of shared marker proteins (**Fig. 2c and Supplementary Fig. 5**) and more distinct protein expression patterns (**Fig. 2d**).

**Fig. 2.**
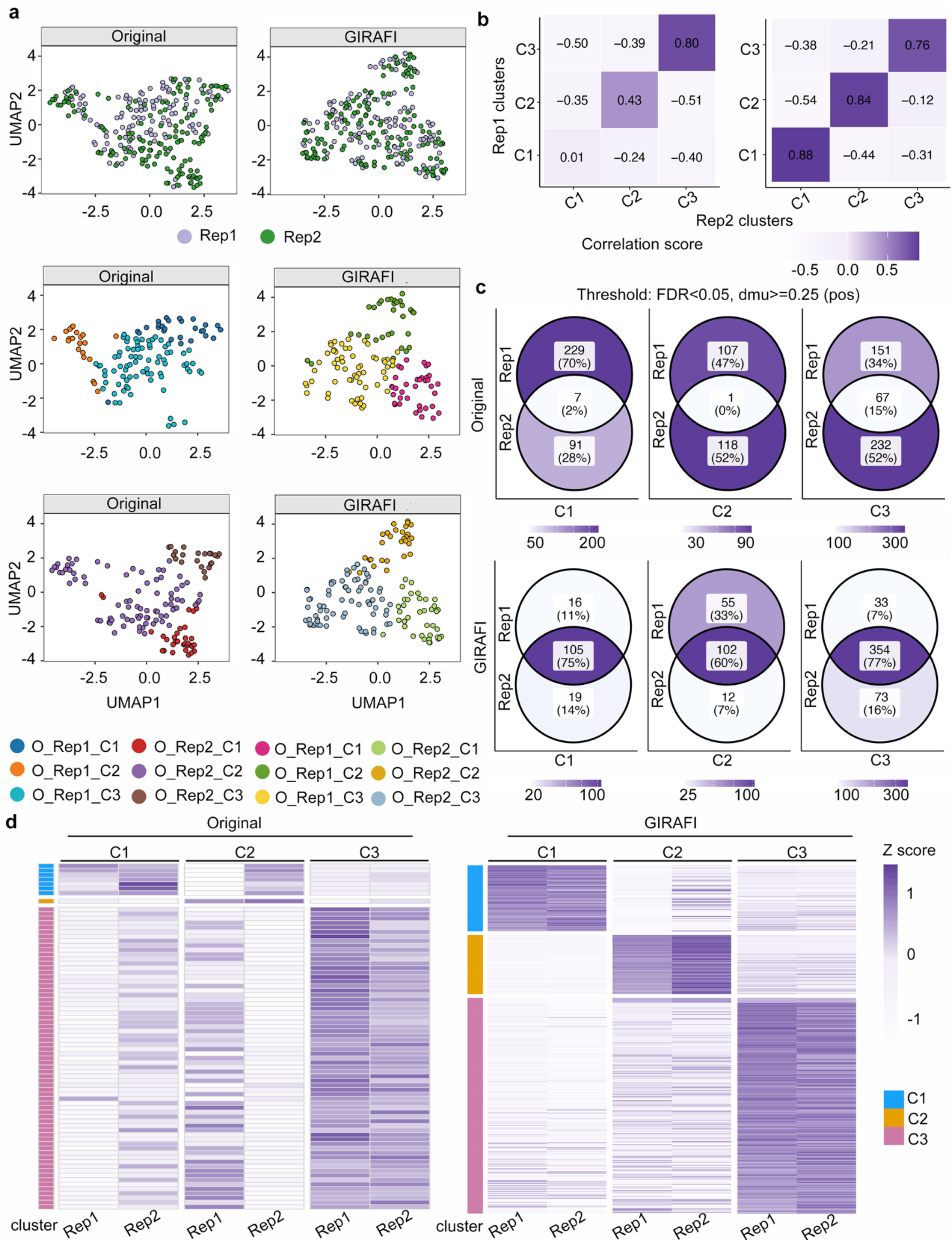
GIRAFI improves cluster structure and cross-replicate concordance in SCP data from *Apis mellifera* forager brains. **a, UMAP comparison between the Original matrix and the GIRAFI matrix**. In the top row, cells are colored by biological replicate (Rep1 and Rep2). In the bottom row, cells are colored by unsupervised cluster labels identified by Single-Cell Consensus Clustering^35^ (SC3; k=3). These cluster assignments were generated separately within Rep1 and Rep2. In the label notation, **O** and **G** indicate the Original and GIRAFI matrices, respectively, and the suffixes indicate the method-replicate-specific cluster identities (For example, O_Rep1_C1-C3 denotes clusters C1 to C3 identified in replicate 1 of the Original matrix). **b, Cross-replicate cluster concordance before and after GIRAFI denoising**. Concordance was quantified as the Spearman correlation between cluster-average protein expression profiles from Rep1 and Rep2. **c, Cluster marker summaries for the three matched clusters**. Markers were defined as proteins significantly enriched in one cluster compared with the other clusters within the same replicate FDR < 0.05, Δ*μ* ≥ 0.25. The top row shows the overlap of marker proteins identified independently in Rep1 and Rep2 from the original data, including the counts and percentages of shared and replicate-specific markers. The bottom row summarizes the marker sets recovered after GIRAFI denoising for each cluster, highlighting the cluster-associated marker composition obtained after imputation. **d, Heatmaps of cluster marker proteins across replicate-by-cluster groups in the Original and GIRAFI matrices**. Columns represent replicate-specific clusters (Rep1-C1-Rep2-C3); rows represent marker proteins grouped by cluster identity, indicated by the lateral color bar. Values denote row-scaled mean expression within each replicate-by-cluster group.

### GIRAFI improves concordance between bulk and pseudobulk protein abundances and recovers known cell types in HEK-derived cell line mixtures

We next benchmarked GIRAFI in a three-component cell-line system consisting of parental Human Embryonic Kidney (HEK) cells (n = 30) and two engineered HEK derivatives expressing T-type calcium channels (hCav3.1, n = 37; hCav3.2, n = 39) (**Supplementary Fig. 6**). Importantly, this benchmark included matched bulk proteomic measurements for the same parental, hCav3.1 and hCav3.2 populations. We therefore used the known population identities to assess single-cell label recovery, and compared pseudobulked single-cell profiles with the matched bulk measurements to evaluate consistency between single-cell and bulk proteomic signals.

To first test whether imputation improves protein-level signal, we compared pseudobulked SCP data to the matched bulk measurements. GIRAFI substantially increased pseudobulk concordance across all three populations (Spearman’s correlation: HEK = 0.60, hCav3.1 = 0.43, and hCav3.2 = 0.43) relative to the Original matrix (Spearman’s correlation: HEK = 0.20, hCav3.1 = 0.06, and hCav3.2 = 0.10) (**Fig. 3a**). Compared to the original data, GIRAFI yielded clearer cell-type-specific protein abundance profiles (**Fig. 3b**; **Supplementary Fig. 7**). A cell-type-specific marker analysis performed separately within each HEK-derived group further showed more complete recovery of expected markers in GIRAFI (**Supplementary Fig. 8**).

**Fig. 3.**
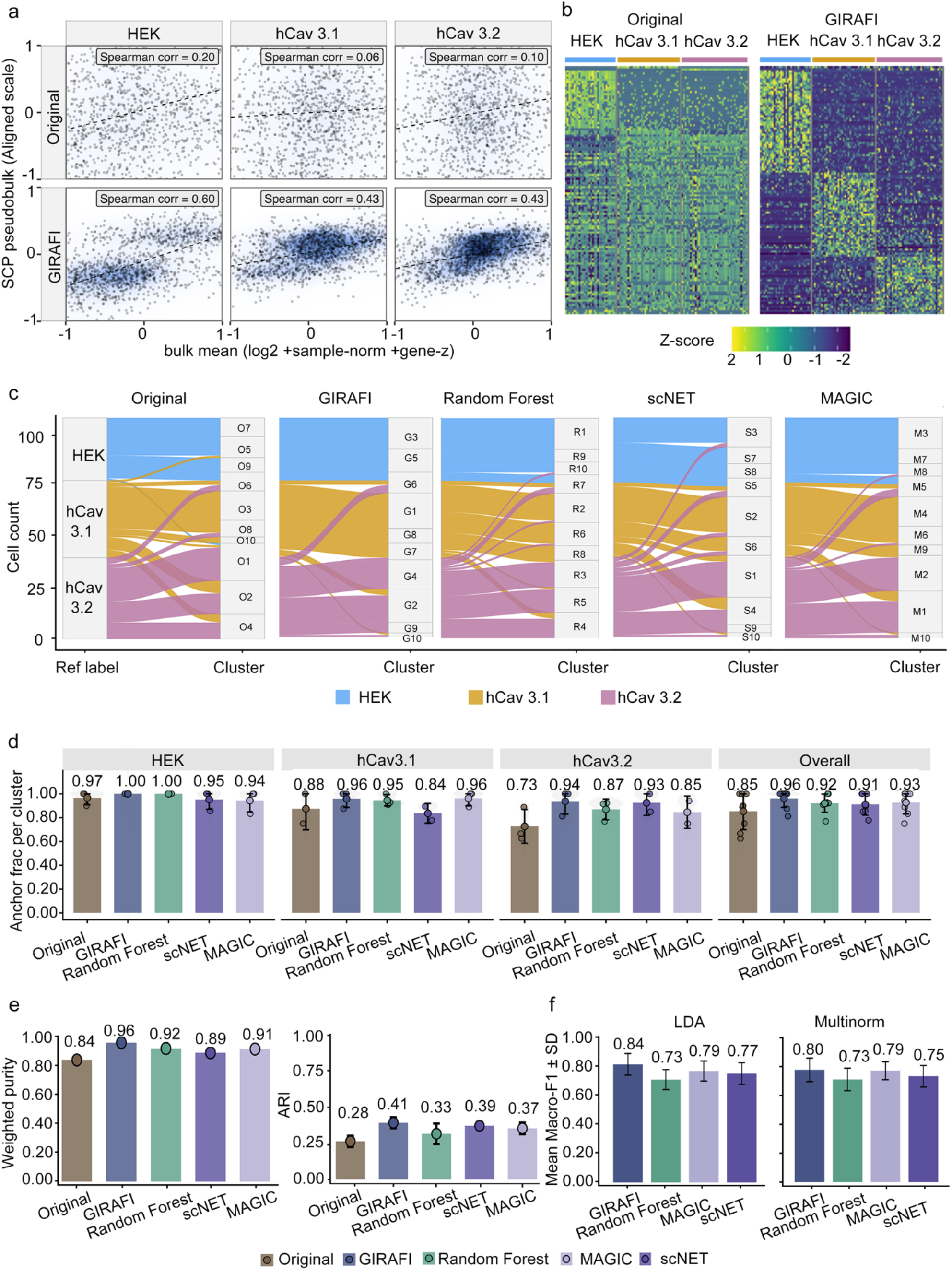
GIRAFI accurately recovers known cell-type composition in an SCP dataset comprising three HEK-derived cell lines and outperforms other state-of-the-art methods. **a, Pseudobulk to bulk proteomics concordance across HEK-derived groups**. HEK-derived groups represent the parental line and two channel-overexpressing derivatives (parental HEK, HEK + hCav3.1, and HEK + hCav3.2). Each point represents one protein. The x-axis shows bulk protein abundance, computed as the mean log_2_ intensity across bulk replicates, followed by sample normalization and per-protein z-scoring. The y-axis shows SCP pseudobulk protein abundance, computed as the mean log_2_ intensity across single cells from the corresponding HEK-derived group and transformed using the same per-protein scaling as the bulk data. Insets report Spearman correlation coefficients. Blue shading indicates local point density, with darker regions representing a higher density of proteins. **b, Heatmaps of cell-by-protein expression before and after GIRAFI denoising**. Expression values are z-scored, and cells are ordered by HEK-derived line (parental HEK, HEK + hCav3.1, and HEK + hCav3.2) in the Original and GIRAFI matrices. **c, Alluvial plots linking reference labels to unsupervised cluster assignments across methods**. Reference labels are shown on the left and unsupervised cluster assignments on the right for each method panel. Flows represent cell counts, and colors indicate reference labels. For visualization and local assignment analysis, cells were further grouped into 10 local neighborhoods. **d, Anchor fraction per cluster across denoising methods**. Anchor fraction is shown for each reference type and overall across the different methods (Original, GIRAFI, Random Forest, scNET, and MAGIC). Here, anchor fraction denotes the fraction of cells within a neighborhood cluster that are anchored to a given reference type in the local neighborhood analysis. Bars show the mean across local neighborhood clusters; error bars indicate SD across clusters. **e, Global clustering agreement metrics across methods**. Clustering performance was quantified using weighted purity and adjusted Rand index (ARI). **f, Five-fold cross-validation performance for downstream cell-line classification across denoising methods**. Classification (**Supplementary Note**) was performed using matrices generated by GIRAFI, Random Forest, scNET, and MAGIC. Results are summarized as mean macro-F1 ± SD. across folds for linear discriminant analysis^36^ (LDA) and multinomial logistic regression^37^ (Multinom).

GIRAFI denoising also improved agreement between known cell-type labels and unsupervised clusters compared to non-imputed data and other imputation methods (**Supplementary Fig. 9**). To visualize the correspondence in local cell-neighborhood space (see **Methods**), we used alluvial plots to compare the three ground-truth HEK-derived groups with unsupervised cell-neighborhood assignments derived from the denoised SCP matrix generated by each imputer (**Fig. 3c**). Consistent with this result, GIRAFI increased the anchor fraction (i.e., the fraction of cells in each cluster assigned to a single reference group), indicating that inferred clusters were more strongly dominated by a single known cell-type label (**Fig. 3d**; see **Methods**). GIRAFI further improved agreement between unsupervised clustering structures and known labels, as reflected by higher weighted purity and increased Adjusted Rand Index (ARI) relative to the Original matrix (**Fig. 3e**).

If imputation captures valid cell-type-specific protein signals, classifiers trained on the denoised matrix should more reliably distinguish the three HEK-derived populations under held-out evaluation. In a five-fold cross-validation cell type classification task, classifiers trained on the denoised matrix generated by GIRAFI achieved the highest cell-type classification performance compared to existing methods for data imputation (**Fig. 3f**).

### GIRAFI enables cell type integration across peripheral blood and bone marrow mononuclear cells

In practice, SCP data are often compared across experiments, tissues, or antibody-enrichment strategies, and are not limited to analyses within a single dataset. A useful denoising method should therefore reduce source-driven batch effects among biologically related populations while maintaining cell type-specific patterns. To test this, we analyzed two SCP datasets collected from distinct sources: FACS-sorted bone marrow mononuclear cells (BMMCs; 311 cells total, including 77 CD13+ monocytes cells, 80 CD4+ T cells, 85 CD8+ T cells, and 69 CD19+ B cells) and magnetically enriched peripheral blood mononuclear cells (PBMCs; 116 cells enriched for monocytes and B cells). These datasets shared overlapping immune populations but differed in tissue source and experimental setup, allowing us to examine if imputation with GIRAFI improved integration of related cell types across datasets.

Before imputation, cells clustered primarily by sample source (BMMC versus PBMC) and not cell type (**Fig. 4a**). After GIRAFI imputation, cells from the shared CD19+/B-cell and CD13+/monocyte-like compartments clustered more closely across the two datasets while remaining distinct from BMMC-specific CD4+ and CD8+ populations (**Fig. 4a**). Dataset-specific clustering metrics supported the same overall trend. After imputation, unsupervised clusters showed preserved or improved agreement with the annotated cell-type labels, as reflected by cluster purity and silhouette scores (**Supplementary Fig. 10 and Supplementary Note**).

**Fig. 4.**
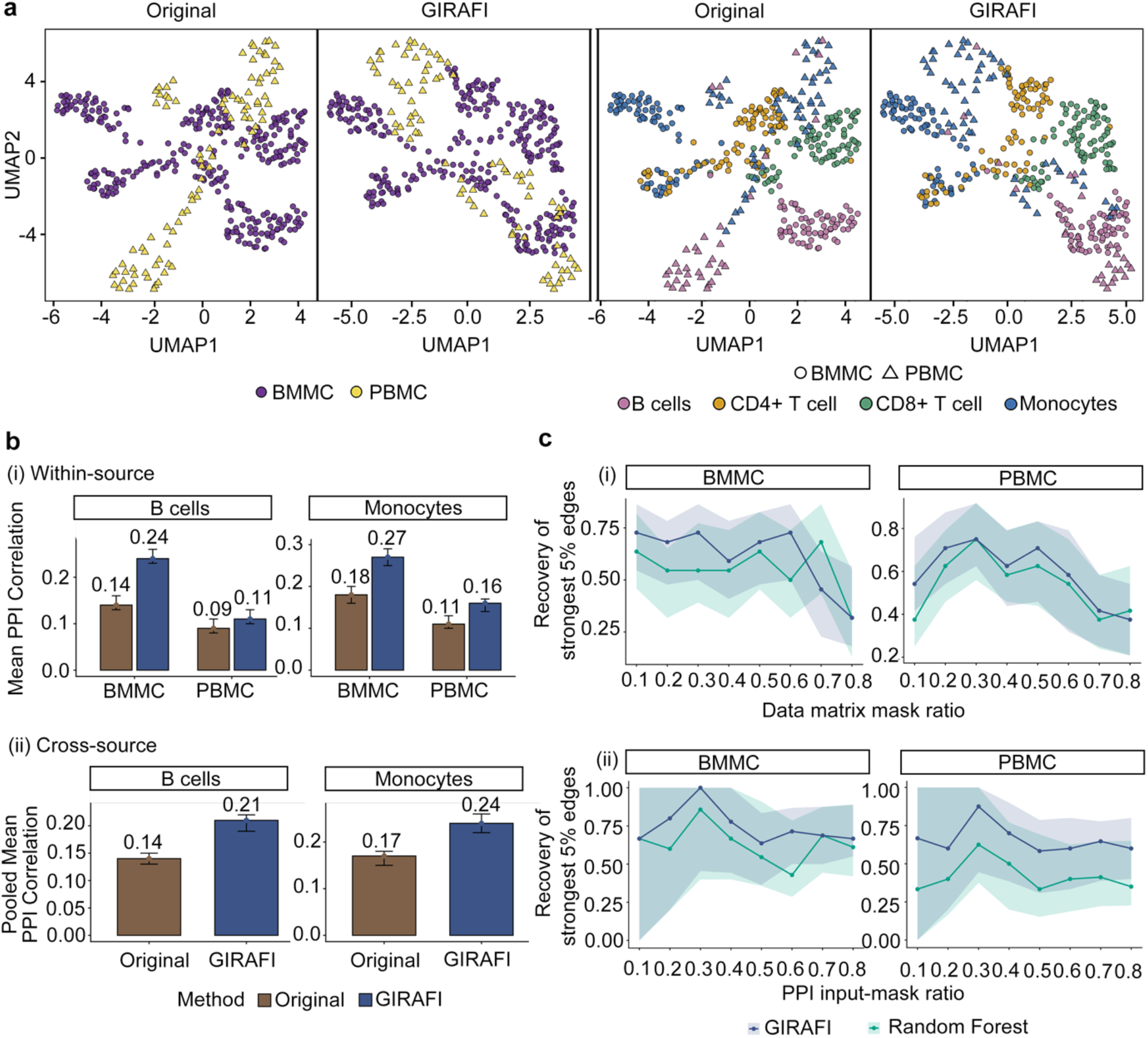
GIRAFI improves cross-source integration and preserves PPI-consistent protein covariance between bone marrow-derived mononuclear cells (BMMCs) and peripheral blood mononuclear cells (PBMCs) proteomes. **a, Joint UMAP embeddings of BMMC and PBMC before and after GIRAFI imputation**. In the left two panels, cells are colored by source, with BMMC shown as purple circles and PBMC as yellow triangles. In the right two panels, the same embeddings are colored by annotated cell type, including B cells, CD4+ T cells, CD8+ T cells, and monocytes, with BMMC shown as circles and PBMC as triangles. **b, Protein-to-neighbor correlation based on the PPI graph. (i)**, Within-source mean protein-to-neighbor correlation for B cells and monocytes in BMMC and PBMC. For each protein, correlations with its PPI neighbors were computed across cells within the indicated source and cell type, and then averaged across neighbors and proteins. Error bars indicate bootstrap 95% confidence intervals across proteins. (**ii)**, Cross-source pooled mean protein-to-neighbor correlation after combining matched cell types across BMMC and PBMC. The same PPI-neighbor correlation metric was computed after pooling cells from matched cell types across sources. Error bars indicate bootstrap 95% confidence intervals across proteins. **c, Recovery of the strongest 5% PPI-consistent edges under masking-based evaluation**. (**i)**, Recovery performance across increasing data-matrix mask ratios. (**ii)**, Recovery performance across increasing PPI input-mask ratios at a fixed 10% data-mask ratio. In both analyses, the strongest 5% PPI-consistent edges were defined from the reference protein-correlation structure and evaluated after masking-based reconstruction. Shaded ribbons indicate 95% confidence intervals.

We next considered if the improved embedding space observed in **Fig. 4a** was accompanied by better recovery of protein covariance along known biological relationships. Using the PPI graph, we quantified the mean protein-to-neighbor correlation within each source and cell type. GIRAFI increased this metric in both B cells and monocytes from both BMMCs and PBMCs, with the largest gains observed in B cells (0.14 to 0.24) and monocytes (0.18 to 0.27) isolated from BMMCs (**Fig. 4b(i)**). When matched cell types were pooled across sources, the same pattern remained, with pooled neighbor correlation increasing from 0.14 to 0.21 in B cells and from 0.17 to 0.24 in monocytes (**Fig. 4b(ii)**).

For the final test, we evaluated robustness using two masking-based recovery analyses (See **Methods**). Performance was assessed on held-out observations that were excluded from model fitting, whereas in the PPI-input-mask analysis, subsets of prior edges were removed before graph construction and then used only for downstream evaluation. Across increasing data-mask and PPI-mask ratios, GIRAFI generally maintained higher recovery than Random Forest, which served here as the strongest baseline imputer in the NRMSE benchmark. The results showed that GIRAFI better preserved the biologically meaningful protein-protein covariance under both observation-level missingness and prior network information incompleteness (**Fig. 4c**).

### Astrocyte infection data provide a complementary test of GIRAFI under donor-specific and perturbation-driven variation

To evaluate the utility of GIRAFI for denoising, we applied GIRAFI to a human astrocyte infection dataset containing 733 single cells from two donors under control and virus-treated conditions. Because donor-to-donor variation can be substantial in primary-cell datasets, we did not assume that infection-associated variation should exceed donor-associated variation. Instead, we evaluated whether imputation preserved infection-associated structure while avoiding embeddings that were dominated by donor- or sparsity-associated patterns. In the original matrix, the astrocyte embedding was driven by donor differences. After GIRAFI denoising, we observed a clearer separation of condition (**Fig. 5a**). At the matrix level, GIRAFI also increased within-group similarity across donor-condition subsets, supporting the improved organization observed in the embedding (**Supplementary Fig. 11**).

**Fig. 5.**
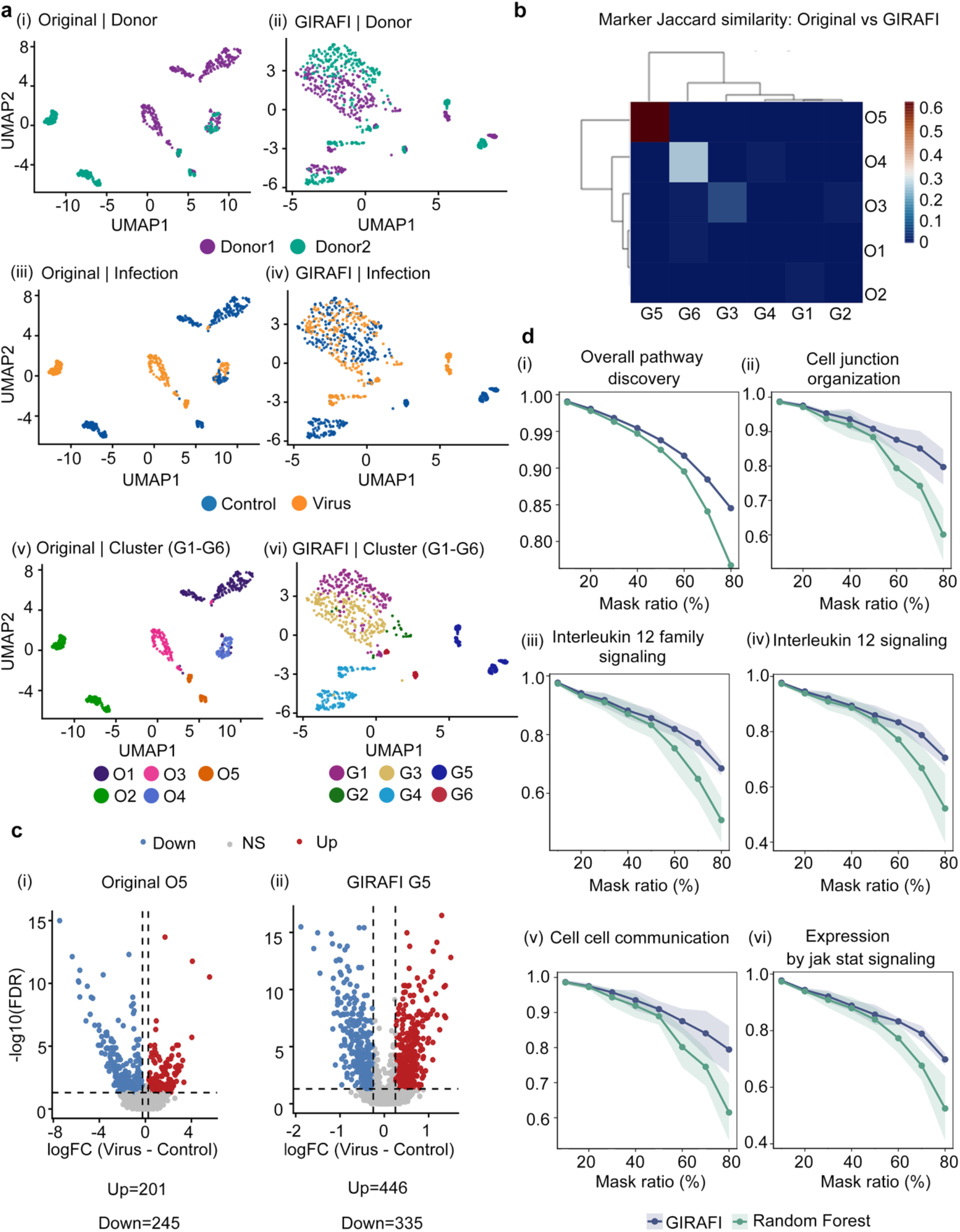
GIRAFI enables cross-condition data comparison and identifies infection-specific cell states in an astrocyte SCP dataset. **a, UMAP embeddings of the astrocyte infection dataset before and after GIRAFI denoising**. Embeddings were generated from the Original and GIRAFI matrices. Panels are colored by donor identity (**i, ii**), donor-condition group (**iii, iv**), and unsupervised cluster assignments (**v, vi**). **b, Marker-set correspondence between Original and GIRAFI clusters**. The heatmap shows Jaccard similarity between cluster marker protein sets, with rows representing Original clusters (O1-O5) and columns representing GIRAFI clusters (G1-G6). Rows and columns are ordered by hierarchical clustering, and higher values indicate greater overlap between the marker programs recovered from the two matrices. **c, Differential protein abundance between virus and control conditions within a representative cellular state**. Volcano plots show within-cluster differential expressions for Original cluster O5 and its GIRAFI counterpart G5, with log_2_ fold change (Virus - Control) on the x-axis and −log_10_ (FDR) on the y-axis. Proteins passing the significance thresholds are colored as increased or decreased in the virus condition, and the total numbers of significantly increased and decreased proteins are annotated in each panel. **d, Pathway-level recovery under increasing missingness**. Starting from the complete subset of the reference matrix, observed entries were randomly masked at rates from 10% to 80%, followed by imputation with GIRAFI or Random Forest and re-estimation of pathway activity. Curves show concordance with the reference pathway signal (y-axis; higher values indicate better recovery) as a function of mask ratio (x-axis), for overall pathway recovery and representative biological programs. Shaded bands indicate variability (95% CI) across runs.

We next assessed if imputation preserved known astrocyte states. Because the Original representation supported a five-cluster solution (**Supplementary Fig. 12**), we used this granularity as the reference for cross-method comparison. Marker-set overlap analysis showed that GIRAFI did not distort the original biological state structure. Some GIRAFI clusters still matched the Original clusters, but GIRAFI also separated one additional subpopulation that was not resolved in the Original matrix (**Fig. 5b**). The same pattern was observed when GIRAFI was re-clustered at fixed k, confirming that the denoised representation retained coherent state relationships under matched granularity (**Supplementary Fig. 13**). Together, these findings indicate that GIRAFI refined the existing astrocyte state structure while preserving its overall organization.

We then examined whether the increased cross-donor overlap reflected attenuation of donor-associated structure, which may include both technical and biological sources of variation, while preserving infection-associated states. Reactome enrichment analysis of the fixed number of clusters showed that virus- and immune-response programs were more clearly concentrated in GIRAFI-resolved states G4 and G5 containing host-virus interaction and cytokine-response terms that were weaker or less distinctly localized in the Original matrix (**Supplementary Fig. 14**). By contrast, clusters that mixed more readily across donors showed limited enrichment for infection-related pathways, consistent with non-infected astrocytes. This pattern suggests that GIRAFI reduces donor-associated fragmentation without erasing biologically meaningful infection responses. A matched-state comparison further supported this interpretation. Relative to Original cluster O5, the corresponding GIRAFI state G5 produced a stronger virus-versus-control signal, with more significant proteins and a cleaner differential expression profile (**Fig. 5c**). Additional matched-state examples showed similar behavior, and the main directions of differential protein abundance were largely preserved between similar clusters (**Supplementary Figs. 15 and 16**). These results suggest that GIRAFI improves sensitivity to infection-associated changes while maintaining the underlying direction of biological variation.

Finally, we used a controlled mask-and-recover benchmark to test whether the improved structure reflected faithful recovery of biological signals. Observed entries from the four donor-condition groups were randomly masked and then imputed using either GIRAFI or Random Forest, the strongest existing baseline in our benchmark. Pathway activity was then re-estimated and compared with the reference signal. Across increasing levels of missingness, GIRAFI retained higher pathway concordance than Random Forest, both globally and for representative infection-related programs (**Fig. 5d**). GIRAFI also better preserved curated PPI edge-level correlation structure under masking (**Supplementary Fig. 17**), while predictive and latent-factor analyses showed stronger recovery of donor- and infection-associated variation (**Supplementary Fig. 18**). In the astrocyte infection system, GIRAFI enabled more reliable cross-condition analysis by reducing donor-associated technical structure and strengthening recovery of infection-responsive proteomics data.

### GIRAFI restores temporal organization in a PC12 differentiation time-course SCP dataset

We next benchmarked GIRAFI using a nerve growth factor (NGF)-induced rat PC12 differentiation SCP time-course dataset, comprising 149 single cells and 3,839 quantified proteins after quality control, sampled across four experimental timepoints: D0 (n = 33), D2 (n = 29), D4 (n = 39), and D6 (n = 48). Quality-control summaries stratified by sampling day showed that GIRAFI increased the effective number of detected proteins per cell and shifted cells toward higher-coverage bins across timepoints, consistent with reduced sparsity across the time course **(Supplementary Fig. 19)**. This dataset examines a different aspect of denoising performance from the previous discrete-group benchmarks. Here, the key question is whether GIRAFI can retain a continuous temporal progression consistent with the measured sampling time.

In the PC12 time-course dataset, GIRAFI arranged cells from adjacent sampling days into a more continuous time-resolved pattern, whereas the Original matrix displayed a more fragmented representation **(Fig. 6a)**. At the matrix level, protein-by-cell heatmaps grouped by sampling day showed the same trend, with a denser and more continuous expression landscape after GIRAFI than in the dropout-dominated Original matrix **(Supplementary Fig. 20)**. To determine whether this improved organization reflected better temporal ordering, we inferred pseudotime from the Original and GIRAFI representations using the same trajectory-inference procedure and compared the inferred ordering with the experimental time labels. GIRAFI markedly increased the agreement between inferred pseudotime and experimental time, with the Spearman correlation increasing from 0.18 in the Original matrix to 0.60 after GIRAFI recovery **(Fig. 6b, left)**. This gain also extended to pairwise ordering fidelity, with the concordance index (C-index) increasing from 0.57 in the Original matrix to 0.77 after GIRAFI recovery **(Fig. 6b, right)**. The bootstrap intervals shown in **Fig. 6b** summarize the uncertainty around these estimates.

**Fig. 6.**
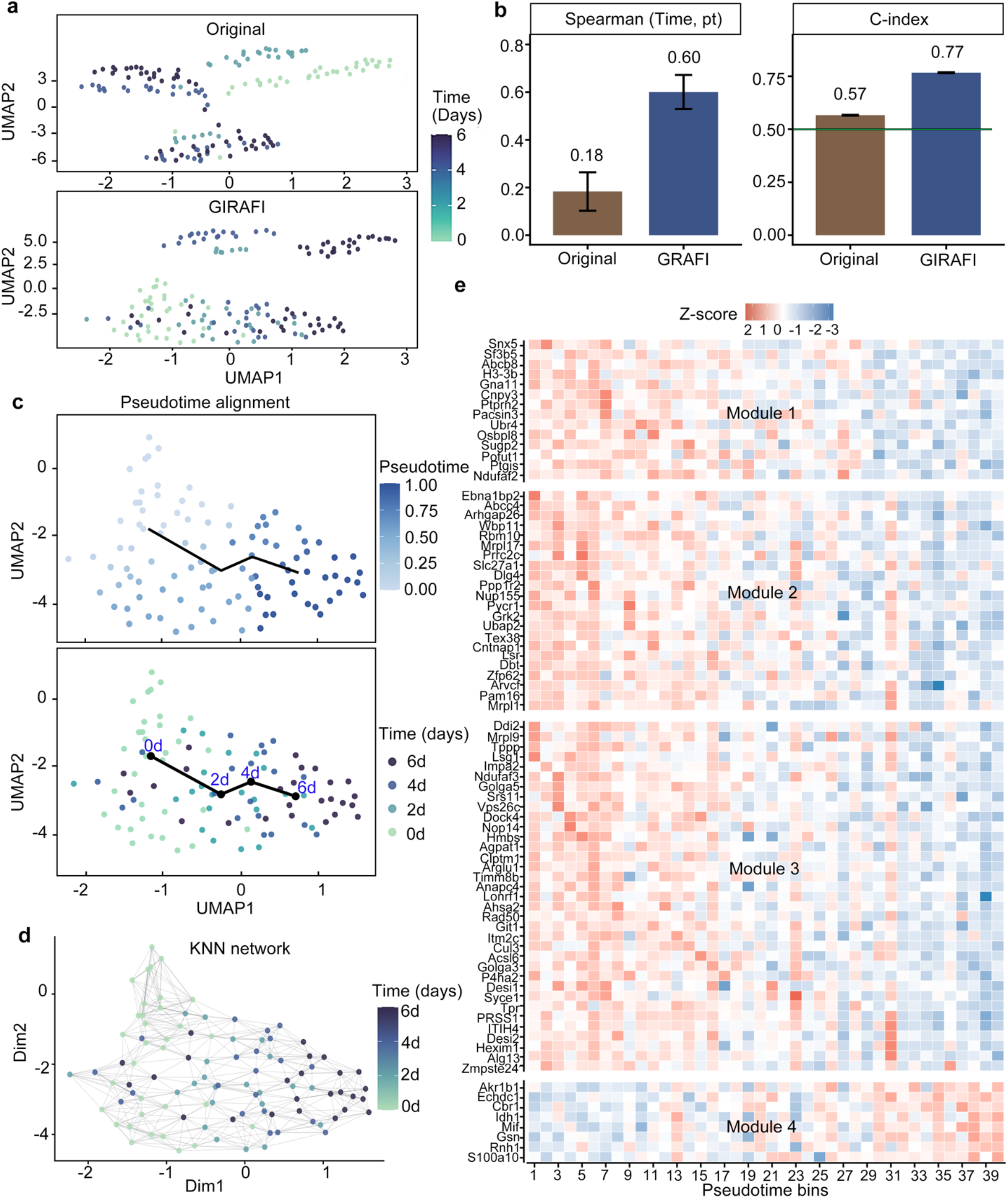
GIRAFI improves trajectory inference accuracy in a PC12 time-course SCP data. **a, UMAP embeddings of the Original and GIRAFI matrices across the PC12 time course**. Two-dimensional UMAP embeddings were computed from the Original matrix (top) and the GIRAFI matrix (bottom). Each point represents a single cell, and colors indicate the experimental sampling time (days). **b, Pseudotime inference accuracy before and after GIRAFI denoising**. Accuracy was quantified by the Spearman correlation between sampling time and inferred pseudotime (left) and by the concordance index (C-index) for pairwise ordering consistency (right). Bars show the mean across bootstrap resamples, and error bars indicate the 2.5th-97.5th percentiles of the bootstrap distribution (95% bootstrap interval). In the C-index panel, the green line marks the random-ordering baseline (C-index = 0.5). **c, GIRAFI centroid-based trajectory projection**. The fitted trajectory curve (black line) summarizes the dominant progression direction through the GIRAFI embedding. In the top panel, cells are colored by inferred pseudotime scaled from 0 to 1. In the bottom panel, the same trajectory is shown with cells colored by discrete sampling time, and timepoint centroids, including D0, D2, D4, and D6, are annotated along the path to show how the inferred trajectory aligns with measured time. **d, GIRAFI-derived k-nearest-neighbor network**. Nodes represent single cells, edges connect nearest neighbors, and node colors indicate experimental sampling day. The network summarizes local cell-cell relationships after GIRAFI recovery and shows how cells from adjacent sampling days are connected along the time-course structure. **e, Heatmap of proteins exhibiting dynamic changes along pseudotime**. Cells are aggregated into pseudotime-ordered bins along the x-axis, arranged from left to right by increasing pseudotime. Rows represent proteins grouped into modules (1-4) based on similar pseudotime-dependent patterns. Values denote row-scaled expression (z-scores), highlighting relative changes along the trajectory within each protein.

To visualize the recovered temporal ordering, we projected a centroid-based trajectory onto the GIRAFI embedding. Pseudotime increased smoothly along the fitted path, and annotated timepoint centroids followed the expected D0-to-D6 progression along the same trajectory **(Fig. 6c)**. The GIRAFI-derived k-nearest-neighbor network further revealed local connections among cells from adjacent sampling days, indicating a continuous time-course organization instead of isolated timepoint-specific clusters **(Fig. 6d)**.

Finally, we evaluated whether the recovered ordering supports interpretable dynamic protein programs. Proteins aggregated into pseudotime-ordered bins formed structured modules with distinct temporal patterns along the trajectory **(Fig. 6e)**. Representative proteins similarly exhibited smooth and significant associations with pseudotime, including increasing, decreasing, and non-monotonic trends **(Supplementary Fig. 21)**. Functional analysis of proteins with increasing intensity along pseudotime further supported the biological relevance of the recovered trajectory, highlighting cytoskeletal and fiber organization, translation-related processes, protein turnover and quality-control pathways, and synapse-related organization terms **(Supplementary Fig. 22)**. The presence of non-monotonic protein trends and module-specific temporal patterns suggests that GIRAFI does not simply impose a uniform smooth gradient across sampling days, but preserves heterogeneous dynamic protein programs along the differentiation trajectory. These results show that, in the PC12 differentiation time-course dataset, GIRAFI restores biologically meaningful temporal organization that is only weakly resolved in the Original matrix.

## Discussion

SCP is advancing rapidly as mass-spectrometry workflows improve, but computational analysis methods remain underdeveloped^2–5^. The data is sparse, incomplete, and shaped by both stochastic sampling and abundance-dependent detection, making recovery difficult to evaluate from reconstruction error alone. A useful SCP imputation method should reduce technical noise while preserving the protein patterns needed for cell-state analysis, marker discovery, pathway interpretation, trajectory inference, and integration across related datasets.

By coupling dataset-aware protein neighborhoods with prior-knowledge-informed graph constraints, we developed GIRAFI as a denoising framework for sparse LC-MS SCP data. Because direct ground-truth measurements remain limited in current SCP workflows, we evaluated GIRAFI by applying it to diverse biological and experimental frameworks. Across distinct biological contexts, GIRAFI improved SCP biological recovery as evaluated by external quantitative agreement, marker detection, pathway analysis, temporal inference, replicate reproducibility, and cross-source integration.

Our results suggest a broader design principle for SCP denoising: prior knowledge is most useful when it guides recovery. GIRAFI leverages prior knowledge by using curated neighborhoods to restrict the candidate predictor space to known interactions, while dataset-specific signals determine which partners are informative in the observed context. Our results indicate that SCP imputation should be framed as biologically constrained signal recovery: the goal is to recover a proteomic framework that remains faithful to the measured context and useful for downstream biological inference. The framework can be extended in several directions. More context-resolved priors, including prior tissue-specific or cell-type-specific PPI maps, protein complex membership, subcellular localization, signaling directionality, and structural constraints, may improve recovery in contexts where active protein neighborhoods differ from generic interaction maps. Integration of proteomics with matched omics layers or other modalities^38^ could provide additional information to distinguish shared regulatory programs from state-specific effects. As SCP datasets increase in size and complexity, scalable implementations and hierarchical modeling strategies will also be needed to apply graph-informed denoising to broader cellular atlases.

Sparse and detection-linked missingness remains a major barrier to robust biological interpretation in LC-MS SCP. GIRAFI addresses this challenge by combining empirical protein associations with curated interaction constraints, improving recovery of protein patterns that support cell-line classification, cross-source integration, pathway analysis, replicate concordance, and trajectory inference. These results support graph-constrained imputation as a practical strategy for improving the reliability of downstream SCP analyses.

## Methods

### Sample collection

#### HEK293

HEK293-F cells (Invitrogen 11625-019) were cultured at 37ºC in a humidified incubator at 5% CO_2_ in Dulbecco’s Modified Eagle Medium (DMEM) supplemented with 10% heat-inactivated FBS (hi-FBS; inactivated for 30 min at 56°C) and 1% non-essential amino acids (NEAAs; GIBCO 11140-050). Stable HEK293-F cell lines expressing the full-length cDNA constructs of the human CaV3.1 (hCav3.1) and human CaV3.2 (hCav3.2) T-type calcium channels were also maintained in the same media with additional 25 µg/ml of Zeocin selection antibiotic (Invitrogen Cat No. R25001). Cells were harvested and dispensed into twin.tec™ PCR Plates 384 well-plates (Eppendorf, Hamburg, Germany) containing 0.5 µL LC-MS graded water using the Uno Single Cell Dispenser (Tecan Trading AG, Switzerland). Samples were stored in the −70°C freezer.

#### Primary astrocytes

Primary astrocytes (ScienCell) from two donors (lot number 31978 and 36204) were cultured in Astrocyte Medium (ScienCell) at 37ºC in a humidified incubator with 5% CO_2_. For each donor, cells were seeded into 6-well plates for both mock control and infection conditions (n = 4 wells per condition) and grown to ~80% confluency. Cells were infected with Human Coronavirus 229E (HCoV-229E) at Multiplicity of Infection (MOI) = 0.2 at 33°C with 5% CO_2_ for 72 hours. Single cells were collected using the same workflow as above.

#### Peripheral blood mononuclear cells

Peripheral blood mononuclear cells (PBMCs) were obtained from StemCell Technologies. To enrich for CD14+ monocytes and CD19+ B cells, PBMCs were stained with an antibody mastermix containing anti-CD14 (61D3) antibodies and anti-CD19 (H1B19) antibodies conjugated to biotin (Biolegend). Magnetic separation was then performed using Dynabeads (Thermo Fisher) according to the manufacturer’s protocol. The flow-through containing monocytes and B cells were dispensed into twin.tecTM PCR 384 well-plates (Eppendorf, Hamburg, Germany).

#### Bone Marrow Mononuclear Cells

Human bone marrow mononuclear cells (BMMCs) were obtained from Lonza (2M-125C, batch 24TL134676) and were stained with a panel of antibodies containing anti-CD13 (WM15), anti-CD19 (HIB19), anti-CD8 (SK1), and anti-CD4 (SK3) antibodies (Biolegend). DAPI-cells were then sorted into twin.tec™ 384 well PCR plates (Eppendorf, Hamburg, Germany) containing 1µL of LC-MS graded water (VWR) using a BD FACSAria Fusion instrument and flash-frozen on dry ice.

#### Honey bee brains

The dissociation protocol was adapted from Zhang et al.^39^. Returning honeybee (*Apis mellifera)* foragers were collected, anesthetized on ice, and dissected under a stereomicroscope to isolate the brain, ensuring removal of salivary and hypopharyngeal glands. The tissue was enzymatically dissociated using 1 mg/mL papain at 37°C for 21 minutes, with mechanical trituration performed every 7 minutes to encourage breakdown. Undigested fragments were removed by centrifugation at 100 x *g* for 15 s, and the supernatant containing the single-cell suspension was collected. Cells were pelleted at 300 x *g* for 5 min, the supernatant was discarded, and the pellet was washed twice with PBS, resuspended, and counted. Single cells were dispensed into twin.tec™ PCR 384-well plates (Eppendorf) containing 1 µL LC-MS grade water using a Tecan Uno Single Cell Dispenser.

#### PC12 cells

PC12 cells (a rat adrenal medulla-derived pheochromocytoma cell line; ATCC CRL-1721) were cultured in Roswell Park Memorial Institute (RPMI) medium supplemented with 5% fetal bovine serum (FBS) (Sigma-Aldrich), 10% horse serum (HS) (Thermo Fisher Scientific), and 1% penicillin-streptomycin (Thermo Fisher Scientific). Cells were differentiated into neuronal-like cells using 100 ng/mL of Nerve Growth Factor (NGF) (Sigma-Aldrich). Following seeding, the medium was replaced with OPTI-MEM (Thermo Fisher Scientific) containing 0.5% FBS to initiate neuronal differentiation. NGF-treated cells were cultured for up to 6 days, with media and NGF replaced every two days to ensure consistent and effective differentiation. Untreated cells were collected before NGF treatment as D0 controls; NGF-treated cells were collected on days 2, 4 and 6 after treatment, washed with PBS (3x) and dispensed into twin.tec™ PCR 384-well plates (Eppendorf) containing 0.5 µL LC-MS grade water using a Tecan Uno Single Cell Dispenser.

### Sample preparation

For each sample, 1 µL of digestion mixture including 1ng/µL Trypsin/LysC in 50 mM Triethylammonium bicarbonate (TEAB) with 0.03% n-dodecyl β-D-maltoside (DDM) was added and incubated in 37°C humidity-controlled incubator for 4 hours. Samples were centrifuged at 2,250 xg for 3min and placed in −70°C freezer for storage.

### LC-MS/MS

Single cell samples were injected and separated on-line using the NanoElute 2 UHPLC system (Bruker Daltonics, Germany) with Aurora Series Gen3 (CSI) analytical columns, (Ion Opticks, Parkville, Victoria, Australia). The analytical column was heated to 50°C using a column toaster M (Bruker Daltonics, Germany). Solvent A consisted of 0.1% aqueous formic acid and 0.5% acetonitrile in water, and solvent B consisted of 0.1% aqueous formic acid and 0.5% water in acetonitrile. Specific LC and MS acquisition parameters are listed below (**Table 1**).

**Table 1.** LC-MS acquisition settings for single-cell proteomics datasets. The table summarizes the chromatographic column configuration, liquid chromatography gradient and flow rate, and mass spectrometer used for each single-cell proteomics dataset analyzed in this study. Column dimensions are reported as length × inner diameter. LC settings indicate flow rate and gradient length.

| Samples | Column type | LC setting | MS |
| --- | --- | --- | --- |
| HEK293/CaV3.1/CaV3.2 | 25 cm × 75 µm i.d. C18 column (1.6 µm particles, 120 Å pores) with CSI fitting | 300nl/min,<br>30min gradient | timsTOF<br>SCP |
| PBMC | 25 cm × 75 µm i.d. C18 column (1.6 µm particles, 120 Å pores) with CSI fitting | 300nl/min,<br>15min gradient | timsTOF<br>SCP |
| BMMC | 8 cm × 75 µm i.d. C18 column (1.6 µm particles, 120 Å pores) with CSI fitting | 450nl/min,<br>7.5min gradient | timsTOF<br>Ultra 2 |
| Astrocytes | 15 cm × 75 µm i.d. C18 column (1.6 µm particles, 120 Å pores) with CSI fitting | 300nl/min,<br>15min gradient | timsTOF<br>SCP |
| PC12 | 25 cm × 75 μm i.d. C18 column (1.6 μm particles, 120 Å pores) with CSI fitting | 300nl/min,<br>30min gradient | timsTOF<br>SCP |
| Honey bee brain | 8cm x 75μm 1.6μm C18 120Å, with CSI fitting | 450nl/min,<br>7.5min gradient | timsTOF<br>Ultra 2 |

### Data analysis

All single-cell data were analyzed using DIA-NN^40^ version 1.8.2 against FASTA databases for single cell samples along with an in-house built common contaminant database. Bulk libraries were generated using 10ng injections and the same method. Single-cell samples were analyzed using a spectral library and the Match-Between-Runs (MBR) feature, leveraging bulk samples for alignment. All parameters used were identical to those previously mentioned^41^.

### Data preprocessing

SCP DIA-NN output tables were processed using a unified pipeline compatible with legacy formats and newer annotation-rich exports. We removed contaminants using a standard contaminant list and excluded entries lacking valid gene symbols. Cells with no detected proteins and proteins with no observed values were discarded. Remaining cells were filtered by total raw intensity and detected-protein counts, and proteins were retained based on a minimum detection fraction (**Supplementary Note**). Intensities were log_2_-transformed using a small pseudocount derived from the minimum positive value, then standardized per protein (z-scoring). Missing values were preserved as NA throughout preprocessing, and proteins with extreme missingness after transformation were removed. All data were processed using R (4.5.2). Single-cell data were processed using the SC3, SingleCellExperiment^42^ package (1.32.0) and Seurat^43^ (4.3.9) in R. Raw (non-imputed) values were used when a zero-preserving reference was needed, while log-scale matrices were used for embeddings and downstream comparisons.

### Imputation baselines

All imputation methods operated on the standardized protein-by-cell matrix with missing entries preserved as NA and were applied to the same filtered feature set for fair comparison. We benchmarked commonly used baselines including cell-based k-nearest neighbors (kNN)^31^, Bayesian PCA (BPCA)^32^, rank-restricted SVD imputation^31^, and iterative random-forest imputation (missForest-style)^33^.

### Protein-protein interaction analysis

To obtain PPI coherence metrics, we used a curated PPI prior downloaded from STRINGdb version 11.5 (ID9606 for human, ID10090 for mouse, and ID7460 for bee)^28^ and computed within-cell-type Pearson correlations for all PPI-connected protein pairs. The fraction of strong PPI edges was defined as the proportion of edges exceeding a fixed positive correlation cutoff within each celltype. The average correlation to PPI neighbors was obtained by averaging correlations from each protein to all of its PPI neighbors, followed by averaging across proteins.

### Graph-informed Random Forest (GIRAFI) imputation

GIRAFI implements two complementary imputation strategies that share the same prior-guided protein graph but differ in how they construct predictors for each target protein. For a given dataset, we run both strategies (GIRAFI-Local, GIRAFI-Neighbor) and select the best-performing variant as the reported GIRAFI result. GIRAFI-Local and GIRAFI-Neighbor were compared using inner-mask NRMSE within the training/masking procedure, and the variant with the lower inner-mask NRMSE was refitted on the corresponding input matrix for downstream analyses.

### Protein association graph construction

#### Association graph

We computed a Spearman correlation matrix using pairwise complete observations, then removed negative correlations and applied an overlap-aware shrinkage based on the number of shared observations:

Let *n* be the number of cells and *n*_*jk*_ be the number of cells where both proteins *j* and *k* are observed.

We formed: 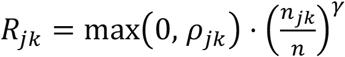, with *γ* = 0.5 by default.

#### Top-k sparsification

For each protein *j*, we kept only the top *K* neighbors (default *K* = 50):

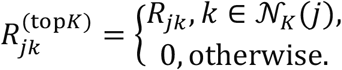

where 𝒩_*k*_ (*j*) denotes the set of top K neighboring proteins of protein *j*.

#### Row-stochastic normalization

We normalized rows to sum to 1 and removed self-loops:

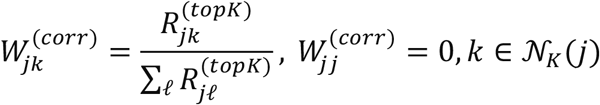

#### Optional PPI graph

If a PPI edge list was provided, we built a weighted adjacency *A*^(ppi)^ and applied symmetric degree normalization:

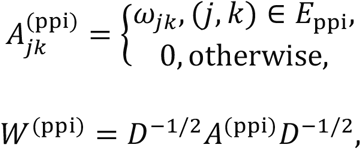

where *D* is the diagonal degree matrix with 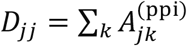.

#### Fusion

The final graph was a convex combination (default *λ* = 0.1):

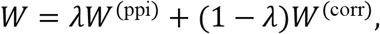

followed by row-stochastic normalization. If no PPI file was available, we used *W* = *W*^(corr)^.

### GIRAFI-Local

For each target protein *j*, we selected the top *K* neighbors 𝒩_*K*_ (*j*) from row *W*_*j*_ (default *K* = 24) and normalized neighbor weights:

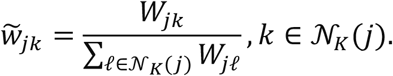

For each cell *i*, we computed neighbor-weighted local statistics using only observed neighboring proteins. Let:

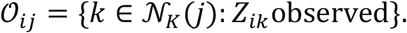

Then,

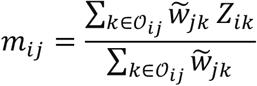

and

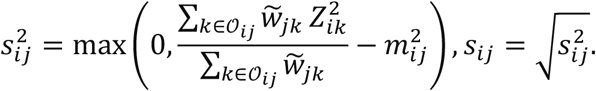

We then trained a random forest regressor for each protein *j*,using (*m*_*ij*_, *s*_*ij*_) as predictors on cells where *Z*_*ij*_ was observed, and predicted *Z*_*ij*_ for cells where it was missing. After this per-protein step, a global random forest refinement pass in a missForest-style was applied to ensure consistency across proteins.

### GIRAFI-Neighbor

For each target protein *j*, we used the neighbor expression vector *Z*_*i*,𝒩(*j*)_ as predictors. Missing values in neighbor predictors were imputed using column medians (within the neighbor submatrix), then a random forest was trained on observed entries of *Z*_*j*_ and used to predict missing entries. A global refinement pass was applied afterward.

### Graph “polish” step

For GIRAFI plans, we applied a light post-processing that modifies only entries that were originally missing. Let 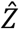 be the imputed matrix and *M* the missingness mask, where *M*_*ij*_ = 1indicates that *Z*_*ij*_ was originally missing. We initialized 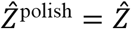 and updated only missing entries as follows:

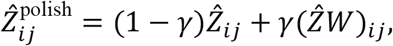

with default *γ* = 0.20 for GraphRF variants. Observed entries were held fixed.

### Mask-and-recover evaluation

To benchmark imputation accuracy, we used a mask-and-recover strategy restricted to a maximal complete (no-NA) submatrix of *Z*. A large NA-free submatrix was identified by a greedy elimination procedure (dropping the “worst” row/column with the most missing entries) with multiple random restarts, subject to minimum size constraints (default ≥ 10 rows and ≥ 50 columns). Within this NA-free submatrix, we masked a fraction *r* of entries (default *r* = 0.10) in a group-balanced manner when group labels were available; otherwise, pseudo-groups were constructed from detected-protein count strata. Performance was quantified on masked entries Ω using:

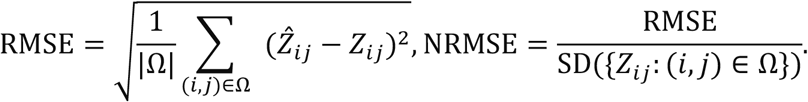

### No-mask imputation outputs for downstream analyses

In addition to masked evaluation, we generated no-mask imputed matrices by running each method directly on the standardized matrix *Z* with its original missingness pattern (i.e., no artificial masking). The resulting completed matrices 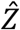 were exported per method and used for downstream analyses (e.g., clustering/visualization/marker analysis/network analysis). Observed (non-missing) entries were preserved; optional graph polishing (when enabled) only updated entries that were originally missing.

### Embedding, clustering, and cluster quality

Dimensionality reduction and visualization by PCA and UMAP were performed using the scater^44^ package (1.38.0). Unsupervised clustering was performed using the package SC3 (1.32.0)^35^, and graph-based clustering was performed using a kNN graph with Louvain community detection. Cluster agreement and separation were evaluated using purity, F1, ARI and silhouette scores and cell label or feature recovery for benchmark samples using precision, recall, and F1-score.

### Markers, statistics, and pathway analysis

Differential testing of protein abundance was performed using Wilcoxon tests with Benjamini-Hochberg false discovery rate (FDR) correction. Effect quantification was performed using within-cluster scoring and, where applicable, area under the curve (AUC). Pathway analyses were performed using ReactomePA^45^ (version 1.48.0) and clusterProfiler^46^ (version 4.12.6). For gene set enrichment analysis (GSEA)^47^, MSigDB^48^ gene sets were retrieved using msigdbr^49^ (version 26.1.0), and enrichment testing was performed using fgsea (version 1.30.0)^50^. Gene annotation was performed using org.Hs.eg.db^51^ (version 3.19.1). Heatmaps were generated using pheatmap^52^ (version 1.0.13), interaction networks were visualized using igraph^53^ (version 2.2.2) and ggraph^54^ (version 2.2.2), and figures were generated using ggplot2^55^ (version 4.0.2) and assembled using patchwork^56^ (version 1.3.2).

### Local cell-neighborhood analysis

To evaluate whether denoising improved cell-type organization at the local cellular level, we constructed a local cell-neighborhood space from each SCP matrix. In contrast to global two-dimensional visualizations such as UMAP^57^, which summarize broad manifold structure, this analysis focuses on local cell-cell relationships encoded by the high-dimensional protein abundance matrix. For each method-generated matrix, cells were represented by their protein abundance profiles, and a nearest-neighbor graph was constructed to identify cells with similar proteomic states.

For each matrix

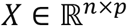

where *n* denotes the number of cells and *p* denotes the number of proteins, rows corresponded to cells and columns corresponded to proteins. Protein abundances were normalized and scaled as described above. Pairwise distances between cells were then computed in the selected feature space, or in a reduced principal-component space when dimensionality reduction was used before neighborhood construction. For two cells *i* and *j*, the cell-cell distance was defined as

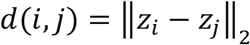

where *z*_*i*_ and *z*_*j*_ denote the protein-profile vectors, or reduced-dimensional embeddings, of cells *i* and *j*. For each cell *i*, its *k* nearest neighbors were identified according to this distance. These relationships were used to construct a cell-cell neighborhood graph

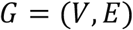

where each node *v*_*i*_ ∈ *V* represents a single cell, and an edge (*i, j*) ∈ *E* indicates that cell *j* belongs to the *k*-nearest-neighbor set of cell *i*, or vice versa after graph symmetrization. Edge weights were optionally defined from cell-cell similarity, for example as an inverse-distance or shared-nearest-neighbor score. This graph defines the local cell-neighborhood space, in which each cell is interpreted according to its nearest proteomic neighbors rather than its global position in an embedding. The resulting graph (**Fig. 3c**) was used as the local cell-neighborhood representation for downstream clustering and label-concordance analysis.

Unsupervised neighborhood assignments were obtained from the cell-neighborhood graph using graph-based clustering. In this framework, a neighborhood cluster represents a group of cells that are locally connected by similar protein abundance profiles. Let

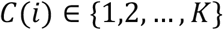

denote the unsupervised neighborhood cluster assigned to cell *i*, and let

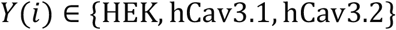

denote the known reference label of the same cell. The correspondence between unsupervised neighborhood assignments and the three known HEK-derived groups was visualized using alluvial plots. In these plots, the left side represents the reference labels *Y*, the right side represents the unsupervised neighborhood clusters *C*, and the width of each flow is proportional to the number of cells shared between a reference group and a neighborhood cluster.

To quantify whether local neighborhoods were dominated by a single known reference group, we calculated an anchor fraction for each neighborhood cluster. For a given neighborhood cluster *c* and reference group *g, n*_*c,g*_ was defined as the number of cells in cluster *c* that belonged to reference group *g*, and *n*_*c*_ was defined as the total number of cells assigned to cluster *c*. The anchor fraction was calculated as

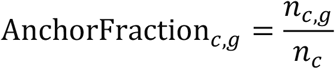

The dominant reference group, or anchor group, of cluster *c* was defined as

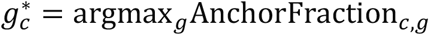

The corresponding dominant anchor fraction was defined as

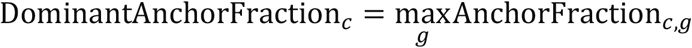

A dominant anchor fraction close to 1 indicates that the neighborhood cluster is almost entirely composed of cells from a single known reference group, whereas a lower value indicates that the neighborhood contains a mixture of multiple HEK-derived populations. Therefore, higher anchor fractions indicate stronger local agreement between the unsupervised cell-neighborhood structure and the known cell-type labels.

To summarize concordance across all neighborhood clusters, we also calculated a weighted purity score:

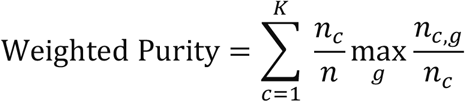

where *n*_*c*_ is the number of cells in neighborhood cluster *c, n*_*c,g*_ is the number of cells in cluster *c* belonging to reference group *g*, and *n* is the total number of cells. This metric gives larger clusters proportionally greater weight and measures the extent to which each unsupervised neighborhood cluster is dominated by a single known group.

In addition, the Adjusted Rand Index (ARI) was used to evaluate the global agreement between the unsupervised neighborhood assignments and the known cell-type labels (**Supplementary Note**). Whereas anchor fraction and weighted purity describe how strongly local neighborhood clusters are dominated by individual reference groups, ARI evaluates the overall pairwise agreement between the two partitions while correcting for chance. Together, these metrics assess whether denoising improves both local neighborhood purity and global consistency with the known HEK-derived cell populations.

### Evaluation of strong edge recovery under data and PPI masking

To evaluate whether each imputation method preserved biologically meaningful protein-protein co-variation structure, we used an edge-recovery metric based on the strongest empirical protein association edges. For each dataset, we first constructed a reference empirical protein association graph from the complete matrix. Edges were ranked by association strength, and the top 5% strongest edges were treated as high-confidence reference edges. After masking and imputation, we reconstructed the empirical protein association graph from each recovered matrix and asked what fraction of the reference strong edges were recovered among the strongest edges in the imputed matrix. We refer to this quantity as the recovery of the strongest 5% edges. Higher values indicate better preservation of high-confidence protein co-variation after imputation.

For the data-masking analysis, increasing fractions of observed values in the data matrix were masked before imputation. Random Forest and GIRAFI were then applied to the masked matrices, and recovery of the strongest 5% reference empirical protein association edges was calculated from the corresponding imputed matrices. To reduce information leakage during GIRAFI model selection, we used a nested masking design. Within each outer-masked dataset, an additional inner mask was generated from the still-observed entries. GIRAFI variants were compared using inner-mask NRMSE, and the variant with the best inner-mask performance was selected. This selected variant was then refitted on the outer-masked matrix and used for the final edge-recovery evaluation.

For the PPI input-masking analysis, increasing fractions of prior PPI edges were withheld from the graph input before GIRAFI imputation. These withheld PPI edges were not used during graph construction or model fitting and were kept only for final evaluation. After imputation, we reconstructed the empirical protein association graph from each recovered matrix and quantified the fraction of withheld PPI edges that appeared among the strongest 5% empirical association edges. This analysis tested whether each method could recover biologically supported protein relationships when the prior interaction graph was incomplete. As above, GIRAFI variant selection was performed using the nested inner-mask procedure, followed by refitting on the outer-masked data before final evaluation. Bootstrap confidence intervals were calculated by resampling edges, and 95% confidence intervals were reported for the edge-recovery curves.

### Latent factor analysis

Comparison of latent structure between non-imputed (Original) and imputed (GIRAFI) matrices generated from single-cell proteomics data of virally infected primary astrocyte cells was performed using MOFA2^58^ (1.14.0). After harmonizing proteins across samples and matching cell-level metadata (donor identity and infection condition), we fit single-view MOFA models (SCP as the only view) separately for non-imputed data or for data imputed using GIRAFI under identical configurations to obtain comparable low-dimensional latent factors. We then quantified how strongly the inferred MOFA2 factors captured donor- and infection-associated variation using factor-covariate association analyses. We complemented these tests with low-dimensional visualizations of the latent space (e.g., PCA/UMAP on factor scores) to assess separation and interpretability across donors and infection conditions.

## Data availability

The datasets generated and/or analyzed during this study will be available in MassIVE under accession number MSV000100666 upon publication. Source data for all main and supplementary figures will be provided with the paper upon publication.

## Code availability

GIRAFI is available through Code Ocean in the peer review process and will be available as open-source software at https://github.com/dorothyzh/GIRAFI upon publication. Reproducible analysis scripts and notebooks to generate all results and figures will be available at the same repository.

## Acknowledgements

We thank Peipei Zhong for assistance with figure preparation; Ziyue Wang for support with the literature review on imputation methodologies; and Lisa Zhang for help with benchmarking imputation performance using public datasets and established imputation approaches; Danning Hao, Qingyang Deng and Zhaowei Chen for helping on the code arrangements. A.E. was partially supported by an Oregon State University Chemistry graduate fellowship and by an HP-OSU collaboration project grant (to C.S.M.). X.T. acknowledge the funding support from Canada Research Chair program (CRC-2024-00164) and the Natural Sciences and Engineering Research Council of Canada (RGPIN-2025-04460).

## Author contributions

H.Z., S. Chi, X.T. and L.J.F. conceptualized the study. S. Chi, R.W., S. Chan, M.L.B., A.E., G.J., J.Y., A.G., T.P.S., C.S.M., M.A.M. and L.J.F. contributed to investigation, data collection and resources. H.Z., S. Chi, Z.W. and J.R. performed formal analysis. H.Z. and Z.W. developed the software and computational workflows. H.Z. developed the mathematical formulation. H.Z., S. Chi and Z.W. prepared the visualizations. X.T. and L.J.F. supervised the study. H.Z. and S. Chi wrote the original draft. All authors reviewed and edited the manuscript.

## References

1. Zhong, H., Chi, S., Magaña, A. A., Fordwour, O. B. & Foster, L. J. Integrative omics and AI-driven systems biology: multilayer networks decoding *Apis mellifera* health and resilience. J. Proteome Res. 24, 5305–5318 (2025).

2. Kelly, R. T. Single-cell proteomics: progress and prospects. Mol. Cell. Proteomics 19, 1739–1748 (2020).

3. Vistain, L. F. & Tay, S. Single-cell proteomics. Trends Biochem. Sci. 46, 661–672 (2021).

4. Guo, T., Steen, J. A. & Mann, M. Mass-spectrometry-based proteomics: from single cells to clinical applications. Nature 638, 901–911 (2025).

5. Cheng, E. et al. SPIN: inkjet-driven nanowell workflow for scalable and sensitive single-cell proteomics. Anal. Chem. 98, 8242–8252 (2026).

6. Shi, Y., Zhong, H., Rogalski, J. C. & Foster, L. J. Optimizing imputation strategies for mass spectrometry-based proteomics considering intensity and missing value rates. Comput. Struct. Biotechnol. J. 27, 1818–1826 (2025).

7. Budnik, B., Levy, E., Harmange, G. & Slavov, N. SCoPE-MS: mass spectrometry of single mammalian cells quantifies proteome heterogeneity during cell differentiation. Genome Biol. 19, 161 (2018).

8. Zhu, Y. et al. Nanodroplet processing platform for deep and quantitative proteome profiling of 10-100 mammalian cells. Nat. Commun. 9, 882 (2018).

9. Petrosius, V. et al. Exploration of cell state heterogeneity using single-cell proteomics through sensitivity-tailored data-independent acquisition. Nat. Commun. 14, 5910 (2023).

10. Gatto, L. et al. Initial recommendations for performing, benchmarking and reporting single-cell proteomics experiments. Nat. Methods 20, 375–386 (2023).

11. Huffman, R. G. et al. Prioritized mass spectrometry increases the depth, sensitivity and data completeness of single-cell proteomics. Nat. Methods 20, 714–722 (2023).

12. Li, S., Li, S., Liu, S. & Ren, Y. Mass spectrometry-based solutions for single-cell proteomics. Genomics Proteomics Bioinformatics 23, qzaf012 (2025).

13. Bantscheff, M., Schirle, M., Sweetman, G., Rick, J. & Kuster, B. Quantitative mass spectrometry in proteomics: a critical review. Anal. Bioanal. Chem. 389, 1017–1031 (2007).

14. Bennett, H. M., Stephenson, W., Rose, C. M. & Darmanis, S. Single-cell proteomics enabled by next-generation sequencing or mass spectrometry. Nat. Methods 20, 363–374 (2023).

15. Vanderaa, C. & Gatto, L. The current state of single-cell proteomics data analysis. Curr. Protoc. 3, e658 (2023).

16. Rosenberger, F. A., Thielert, M. & Mann, M. Making single-cell proteomics biologically relevant. Nat. Methods 20, 320–323 (2023)

17. Yu, S.-H. et al. Quantification quality control emerges as a crucial factor to enhance single-cell proteomics data analysis. Mol. Cell. Proteomics 23, 100768 (2024).

18. Luecken, M. D. & Theis, F. J. Current best practices in single-cell RNA-seq analysis: a tutorial. Mol. Syst. Biol. 15, e8746 (2019).

19. Hou, W., Ji, Z., Ji, H. & Hicks, S. C. A systematic evaluation of single-cell RNA-sequencing imputation methods. Genome Biol. 21, 218 (2020).

20. Svensson, V. Reply to: UMI or not UMI, that is the question for scRNA-seq zero-inflation. Nat. Biotechnol. 39, 160 (2021).

21. Sheinin, R., Sharan, R. & Madi, A. scNET: learning context-specific gene and cell embeddings by integrating single-cell gene expression data with protein-protein interactions. Nat. Methods 22, 708–716 (2025).

22. Eraslan, G., Simon, L. M., Mircea, M., Mueller, N. S. & Theis, F. J. Single-cell RNA-seq denoising using a deep count autoencoder. Nat. Commun. 10, 390 (2019).

23. van Dijk, D. et al. Recovering gene interactions from single-cell data using data diffusion. Cell 174, 716-729.e27 (2018).

24. Hafemeister, C. & Satija, R. Normalization and variance stabilization of single-cell RNA-seq data using regularized negative binomial regression. Genome Biol. 20, 296 (2019).

25. Lopez, R., Regier, J., Cole, M. B., Jordan, M. I. & Yosef, N. Deep generative modeling for single-cell transcriptomics. Nat. Methods 15, 1053–1058 (2018).

26. Love, M. I., Huber, W. & Anders, S. Moderated estimation of fold change and dispersion for RNA-seq data with DESeq2. Genome Biol. 15, 550 (2014).

27. Risso, D., Perraudeau, F., Gribkova, S., Dudoit, S. & Vert, J.-P. A general and flexible method for signal extraction from single-cell RNA-seq data. Nat. Commun. 9, 284 (2018).

28. Szklarczyk, D. et al. The STRING database in 2023: protein-protein association networks and functional enrichment analyses for any sequenced genome of interest. Nucleic Acids Res. 51, D638–D646 (2023).

29. Tsitsiridis, G. et al. CORUM: the comprehensive resource of mammalian protein complexes-2022. Nucleic Acids Res. 51, D539–D545 (2023).

30. Milacic, M. et al. The Reactome Pathway Knowledgebase 2024. Nucleic Acids Res. 52, D672–D678 (2024).

31. Troyanskaya, O. et al. Missing value estimation methods for DNA microarrays. Bioinformatics 17, 520–525 (2001).

32. Oba, S. et al. A Bayesian missing value estimation method for gene expression profile data. Bioinformatics 19, 2088–2096 (2003).

33. Stekhoven, D. J. & Bühlmann, P. MissForest: nonparametric missing value imputation for mixed-type data. Bioinformatics 28, 112–118 (2012).

34. Shen, M. et al. Comparative assessment and novel strategy on methods for imputing proteomics data. Sci. Rep. 12, 1067 (2022).

35. Kiselev, V. Y. et al. SC3: consensus clustering of single-cell RNA-seq data. Nat. Methods 14, 483–486 (2017).

36. Fisher, R. A. The use of multiple measurements in taxonomic problems. Ann. Eugen. 7, 179–188 (1936).

37. Theil, H. A multinomial extension of the linear logit model. Int. Econ. Rev. 10, 251–259 (1969).

38. Zhong, H., Shi, Y., Kozlova, A. et al. Omics insights into the effects of highbush blueberry and cranberry crop agroecosystems on honey bee health and physiology. Proteomics e70033 (2025).

39. Zhang, W. et al. Single-cell transcriptomic analysis of honeybee brains identifies vitellogenin as caste differentiation-related factor. iScience 25, 104643 (2022).

40. Demichev, V., Messner, C. B., Vernardis, S. I., Lilley, K. S. & Ralser, M. DIA-NN: neural networks and interference correction enable deep proteome coverage in high throughput. Nat. Methods 17, 41–44 (2020).

41. Chi, S. et al. Bridging simplicity and depth in single-cell proteomics: a cost-effective workflow and expanded framework for data evaluation. Preprint at 10.64898/2026.02.08.700933 (2026).

42. Amezquita, R. A. et al. Orchestrating single-cell analysis with Bioconductor. Nat. Methods 17, 137–145 (2020).

43. Satija, R., Farrell, J. A., Gennert, D., Schier, A. F. & Regev, A. Spatial reconstruction of single-cell gene expression data. Nat. Biotechnol. 33, 495–502 (2015).

44. McCarthy, D. J., Campbell, K. R., Lun, A. T. L. & Wills, Q. F. scater: pre-processing, quality control, normalization and visualization of single-cell RNA-seq data in R. Bioinformatics 33, 1179–1186 (2017).

45. Yu, G. & He, Q.-Y. ReactomePA: an R/Bioconductor package for Reactome pathway analysis and visualization. Mol. Biosyst. 12, 477–479 (2016).

46. Yu, G., Wang, L.-G., Han, Y. & He, Q.-Y. clusterProfiler: an R package for comparing biological themes among gene clusters. OMICS 16, 284–287 (2012).

47. Subramanian, A. et al. Gene set enrichment analysis: a knowledge-based approach for interpreting genome-wide expression profiles. Proc. Natl. Acad. Sci. USA 102, 15545–15550 (2005).

48. Liberzon, A. et al. Molecular signatures database (MSigDB) 3.0. Bioinformatics 27, 1739–1740 (2011).

49. Dolgalev, I. msigdbr: MSigDB gene sets for multiple organisms in a tidy data format. CRAN v26.1.0, 10.32614/CRAN.package.msigdbr (2026).

50. Korotkevich, G. et al. Fast gene set enrichment analysis. Preprint at 10.1101/060012 (2016).

51. Carlson, M. org.Hs.eg.db: genome-wide annotation for human. Bioconductor R package v3.19.1, 10.18129/B9.bioc.org.Hs.eg.db (2024).

52. Kolde, R. pheatmap: pretty heatmaps. CRAN v1.0.13, 10.32614/CRAN.package.pheatmap (2025).

53. Csardi, G. & Nepusz, T. The igraph software package for complex network research. InterJournal, Complex Systems 1695 (2006).

54. Pedersen, T. L. ggraph: an implementation of grammar of graphics for graphs and networks. CRAN v2.2.2, 10.32614/CRAN.package.ggraph (2025).

55. Wickham, H. ggplot2: Elegant Graphics for Data Analysis (Springer, 2009).

56. Pedersen, T. L. patchwork: the composer of plots. CRAN v1.3.2, 10.32614/CRAN.package.patchwork (2025).

57. McInnes, L., Healy, J. & Melville, J. UMAP: uniform manifold approximation and projection for dimension reduction. Preprint at 10.48550/arXiv.1802.03426 (2018).

58. Argelaguet, R. et al. MOFA+: a statistical framework for comprehensive integration of multi-modal single-cell data. Genome Biol. 21, 111 (2020).

