## Supplementary Information for "Recovering biological structure in sparse single-cell proteomics with GIRAFI"

**Zhong et al.**

**Supplementary Figures**

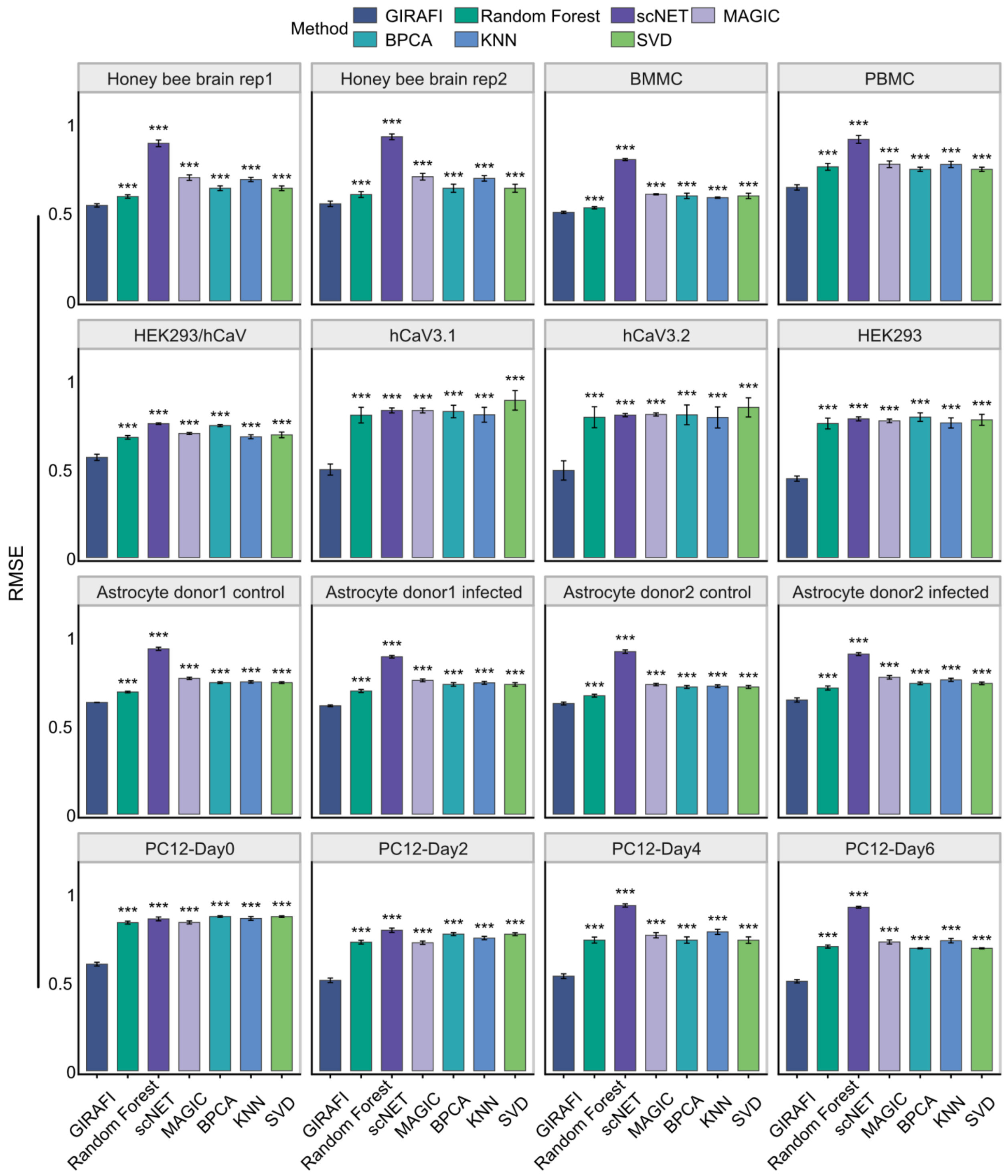

**Supplementary Fig. 1: Imputation performance across datasets assessed by RMSE.**

Bar plots show root mean square error (RMSE) for seven imputation methods across diverse datasets, spanning human samples, established model systems, and less-characterized non-model organisms. Lower RMSE indicates better agreement between imputed and masked true values. Methods include GIRAFI, RF, scNET, MAGIC, BPCA, kNN, and SVD. Error bars represent variability across repeated evaluations. Statistical comparisons were performed using two-sided Welch's *t*-tests with GIRAFI as the reference method; significance is indicated by brackets ($p < 0.05$, $p < 0.01$, $p < 0.001$).

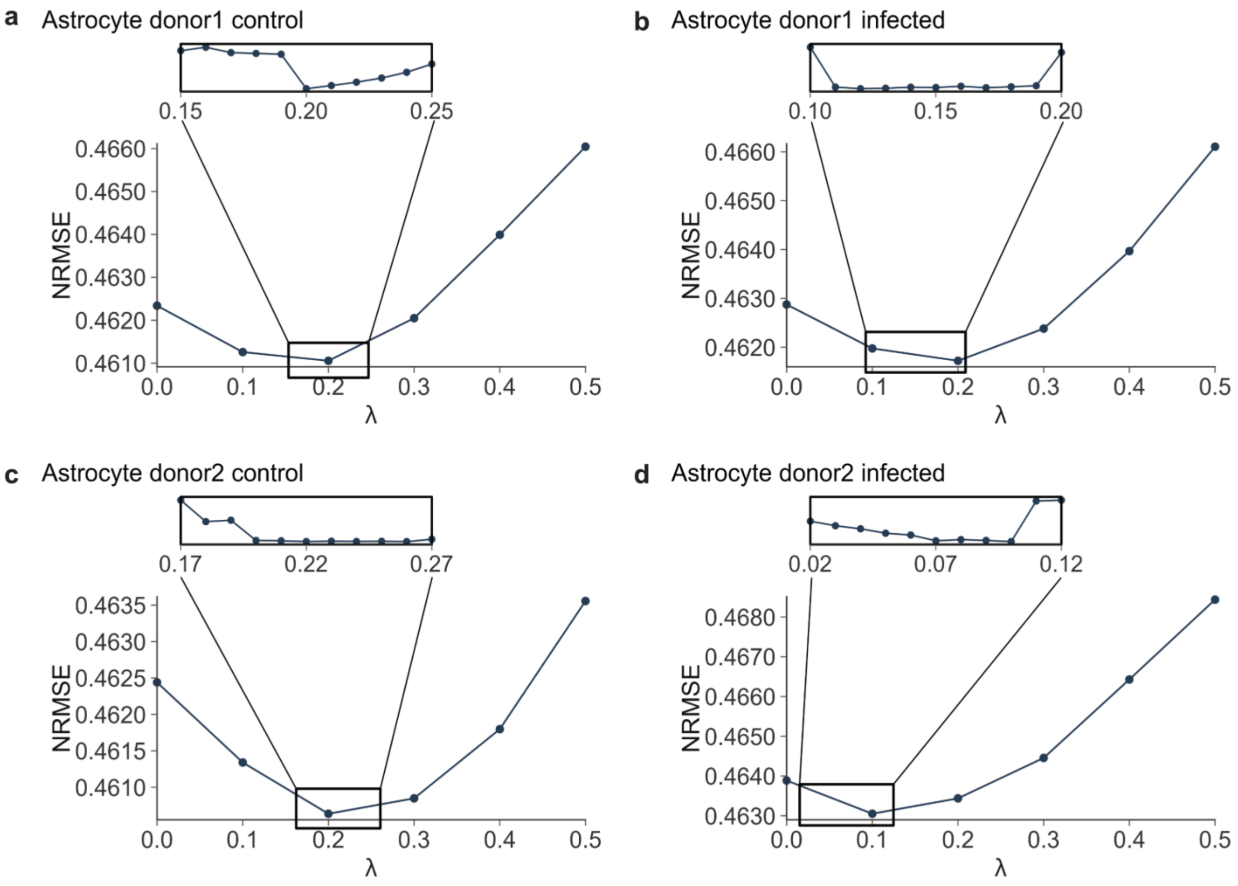

**Supplementary Fig. 2: Tuning of the graph-fusion parameter λ in astrocyte donor- and condition-specific datasets.**

Mask-and-recover performance is measured by NRMSE and shown across candidate values of the graph-fusion parameter λ for four astrocyte subsets: donor 1 control (**a**), donor 1 infected (**b**), donor 2 control (**c**), and donor 2 infected (**d**). Lower NRMSE indicates better recovery performance. In each panel, points represent the mean NRMSE at the corresponding λ value.

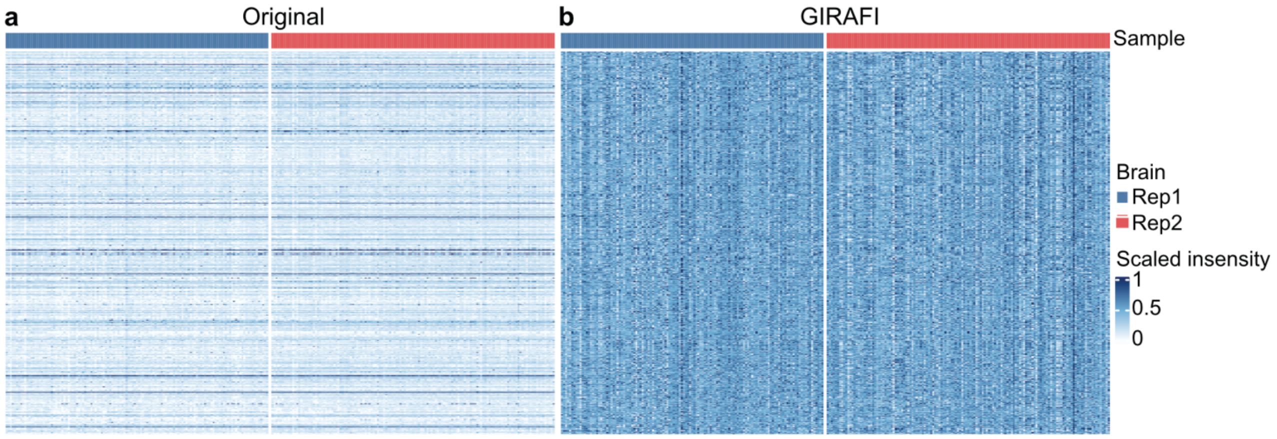

**Supplementary Fig. 3: GIRAFI increases matrix completeness and reduces dropout-driven striping across replicates.**

Heat maps show the protein-by-cell expression matrices for the two biological replicates. **a**, Original matrix before imputation, with zeros fixed and non-zero entries scaled to 0-1. **b**, GIRAFI-denoised matrix, scaled to 0-1. Columns are grouped by replicate, highlighting replicate-wise sparsity in the original data and abundance patterns recovered after GIRAFI denoising. The reproducibility of some dropout patterns across replicates suggests that missingness in SCP is not completely stochastic and may in part reflect non-random processes related to peptide or protein detectability.

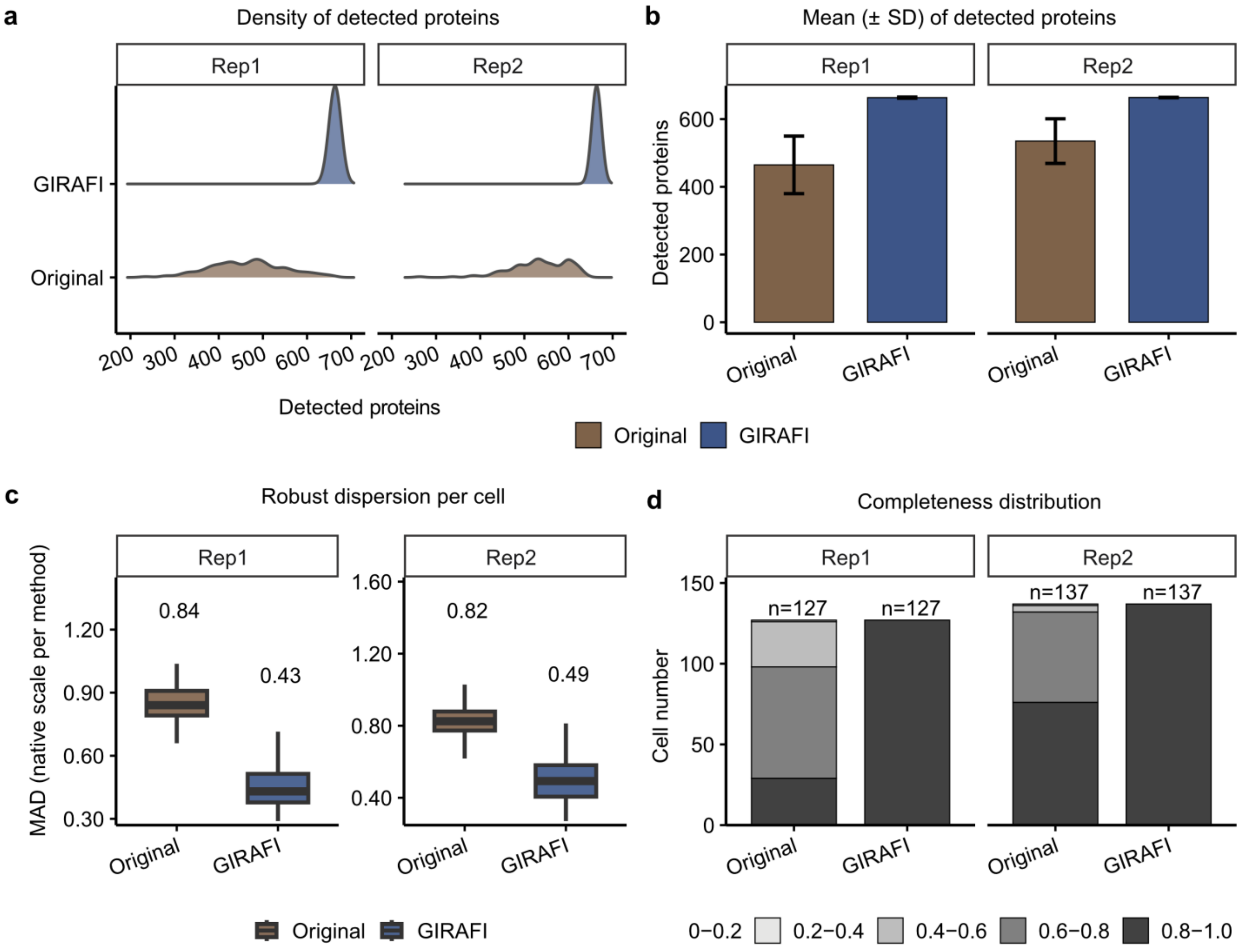

**Supplementary Fig. 4 Quality-control summary for the non-model system highlights improved detection with GIRAFI.**

**a,** Distributions of detected protein features per cell, defined as the number of proteins with non-missing values.

**b,** Mean ± SD of detected protein features per cell across replicates.

**c,** Robust per-cell dispersion. Per-cell dispersion of protein abundances across proteins within each cell after median absolute deviation (MAD)-based outlier removal.

**d,** Completeness distribution across cells. Stacked bars show the number of cells in each per-cell completeness bin, defined by the fraction of observed protein features.

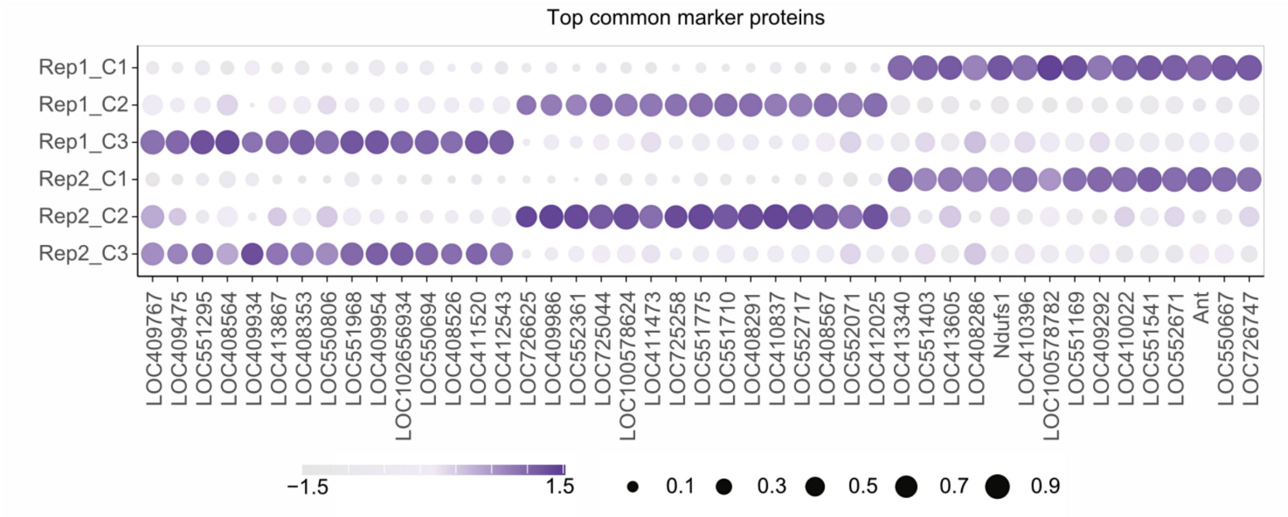

**Supplementary Fig. 5: Common marker proteins show replicate-consistent cluster patterns after GIRAFI.**

Dot plot summarizing common marker proteins across replicates by cluster; dot size indicates detection rate (>0) and color indicates row-scaled mean expression.

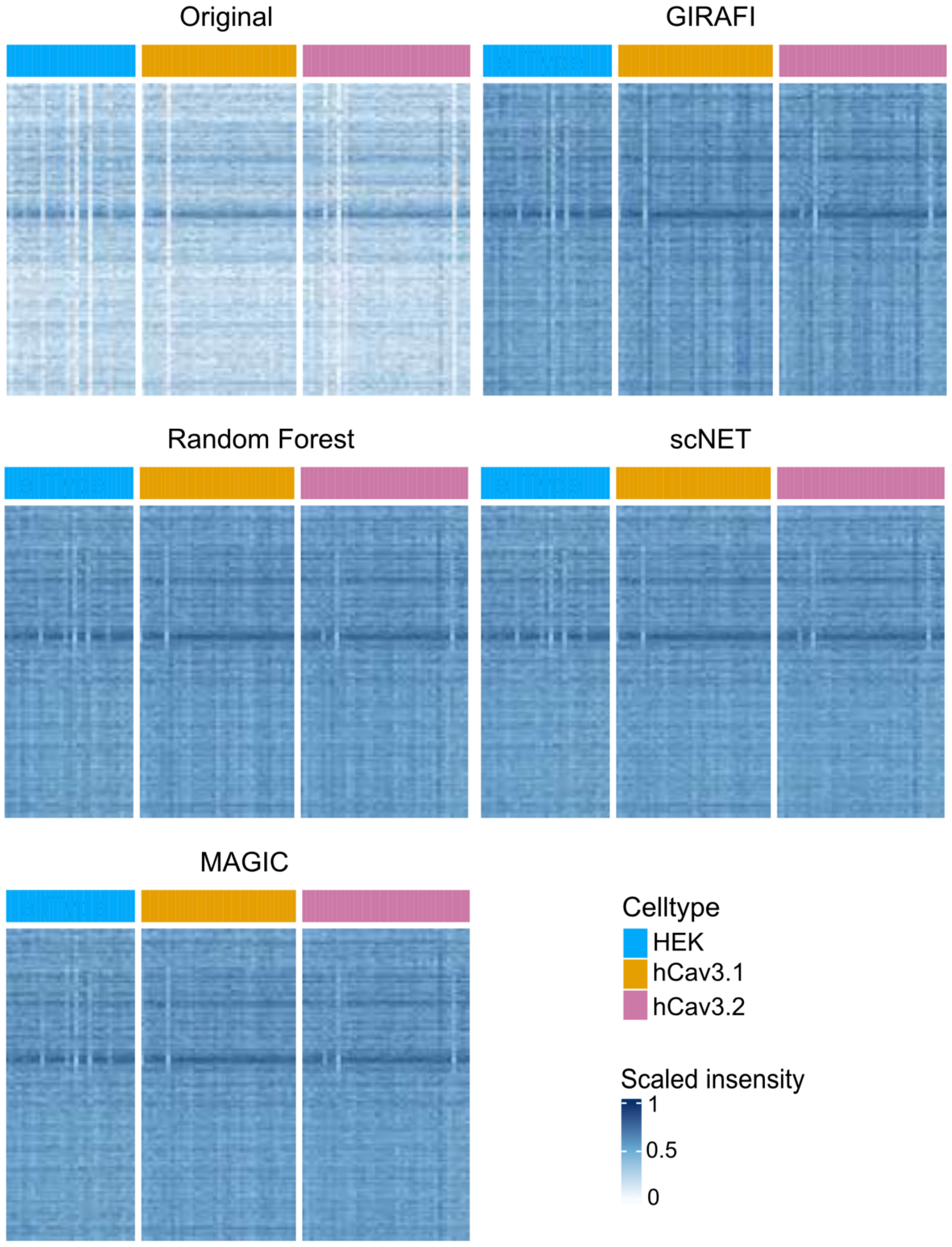

**Supplementary Fig. 6: Matrix-level sparsity patterns across imputers in the HEK/hCav3 mixture**

A protein-by-cell heatmap comparing Original and multiple imputers (GIRAFI, Random Forest, scNET, MAGIC). Values are scaled to 0-1 within each method for visualization, and cells are ordered by reference type (HEK, hCav3.1, hCav3.2); the top annotation indicates cell type.

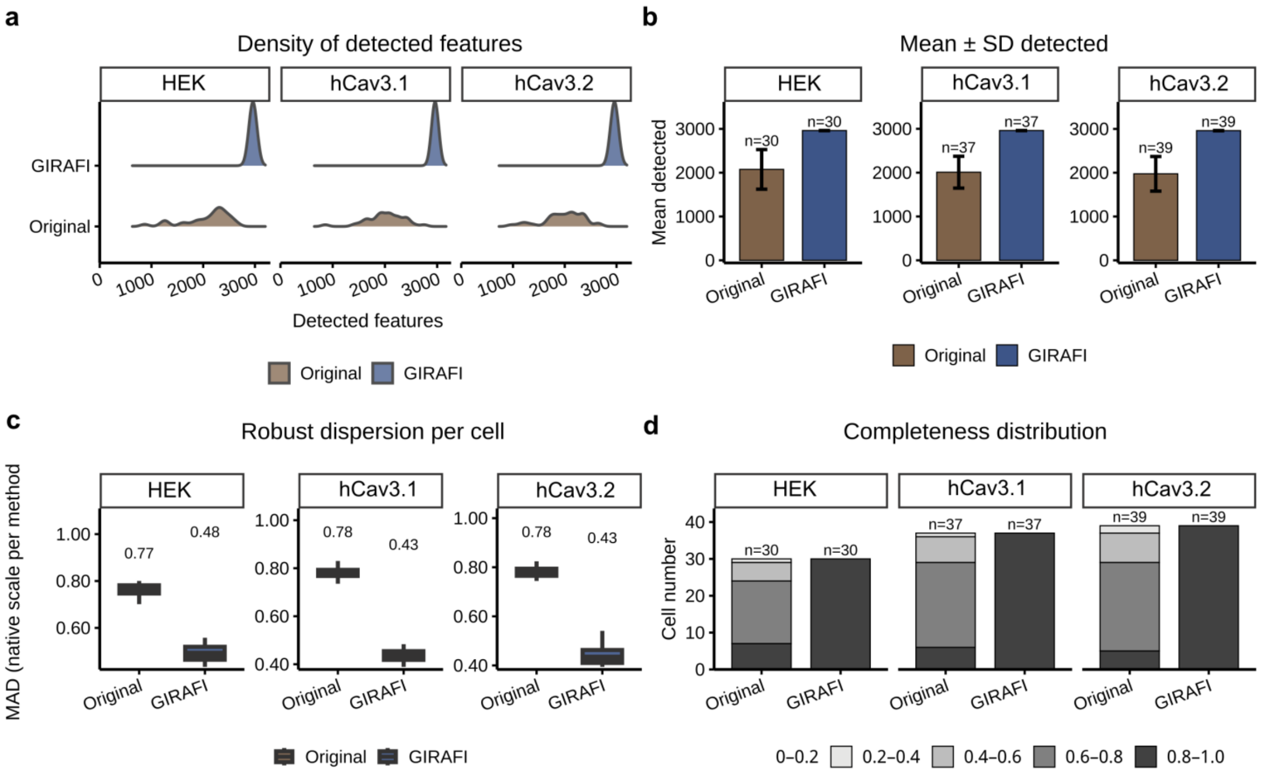

**Supplementary Fig. 7: QC summary by cell type for Original versus GIRAFI**

**a, Density of detected features across HEK-derived groups.** Distributions of per-cell detected proteins, defined as the number of protein values greater than 0, are shown separately for parental HEK, HEK + hCav3.1, and HEK + hCav3.2, comparing the Original and GIRAFI matrices.

**b, Mean ± SD of detected proteins per cell.** Bars summarize the mean number of detected proteins per cell for each HEK-derived group, and error bars indicate the standard deviation.

**c, Robust per-cell dispersion across HEK-derived groups.** Median absolute deviation (MAD), calculated after outlier removal, summarizes within-cell expression dispersion. The Original matrix was computed from $\log_2$(raw intensity) values without a pseudocount, whereas GIRAFI was evaluated using the denoised output on its native scale.

**d, Completeness distributions across HEK-derived groups.** Stacked bars show the number of cells falling into different per-cell completeness bins, defined by the fraction of observed proteins, within each cell type.

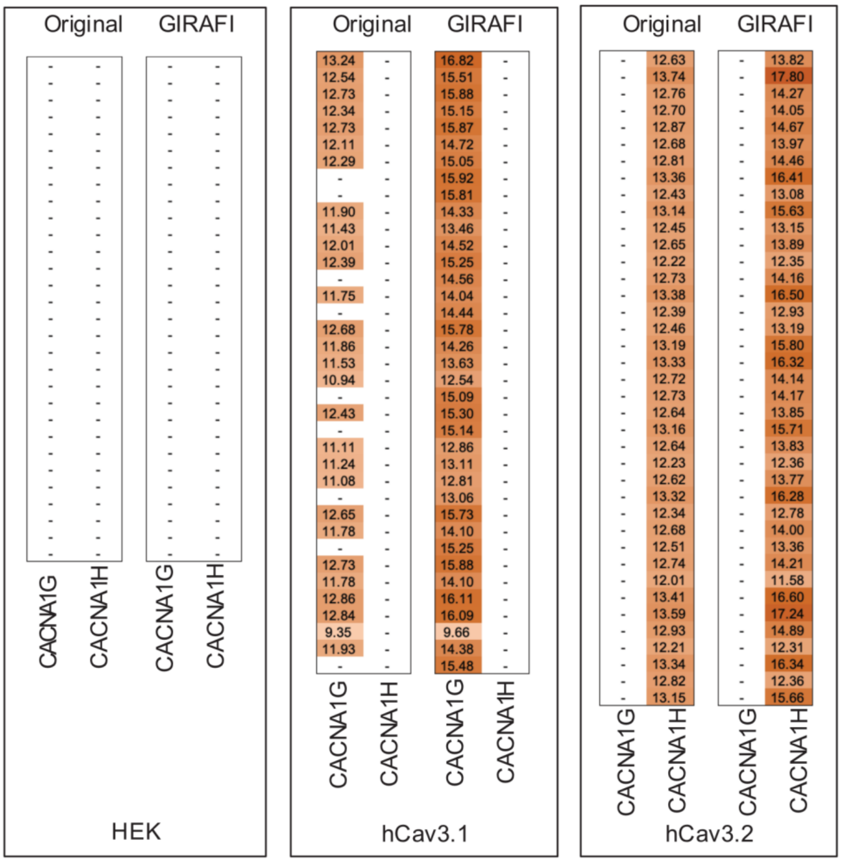

**Supplementary Fig. 8: Recovery of low-detectability channel markers (CACNA1G/CACNA1H)**

For each HEK-derived line type (HEK, hCav3.1, hCav3.2), per-cell signals for channel marker proteins (CACNA1G and CACNA1H) are shown for Original and GIRAFI. Each column corresponds to a marker and rows correspond to single cells (or cell-ordered strips); intensity indicates recovered signal, highlighting differences in detectability and recovery across methods.

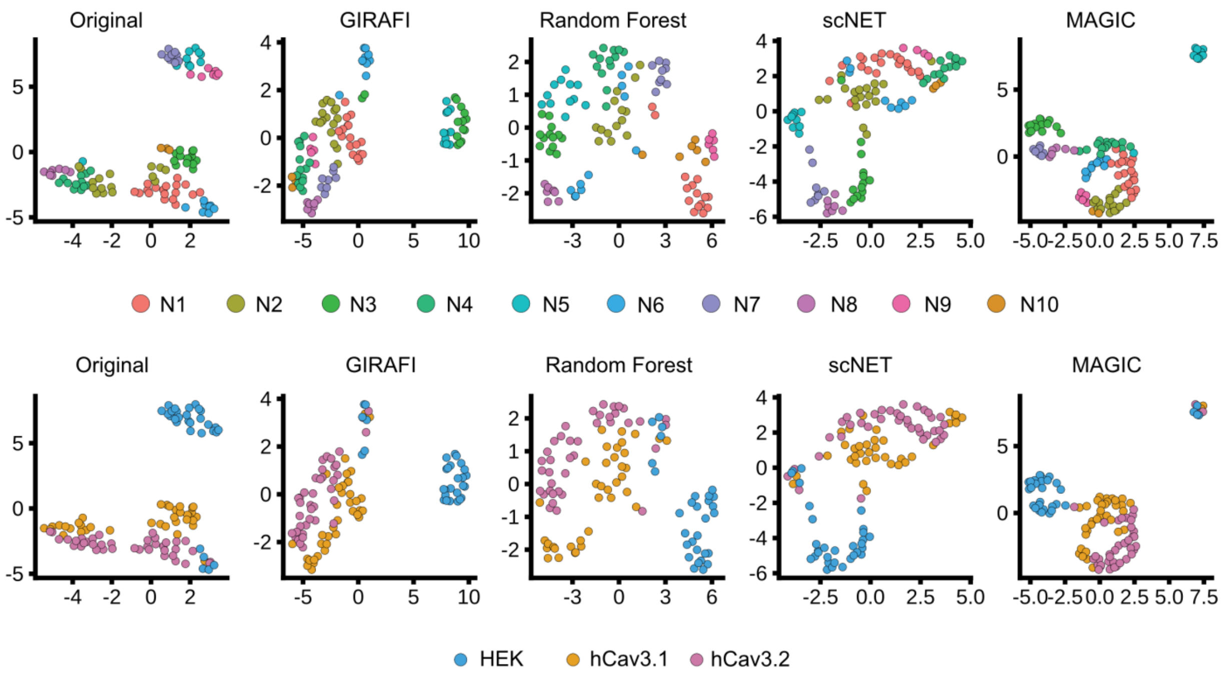

**Supplementary Fig. 9: Neighborhood structure and label coherence in the embedding across methods**

Top row: embeddings colored by neighborhood/graph structure (cluster/community labels) for Original, GIRAFI, Random Forest, scNET and MAGIC. Bottom row: the same embeddings colored by known reference labels (HEK, hCav3.1, hCav3.2) to assess label coherence.

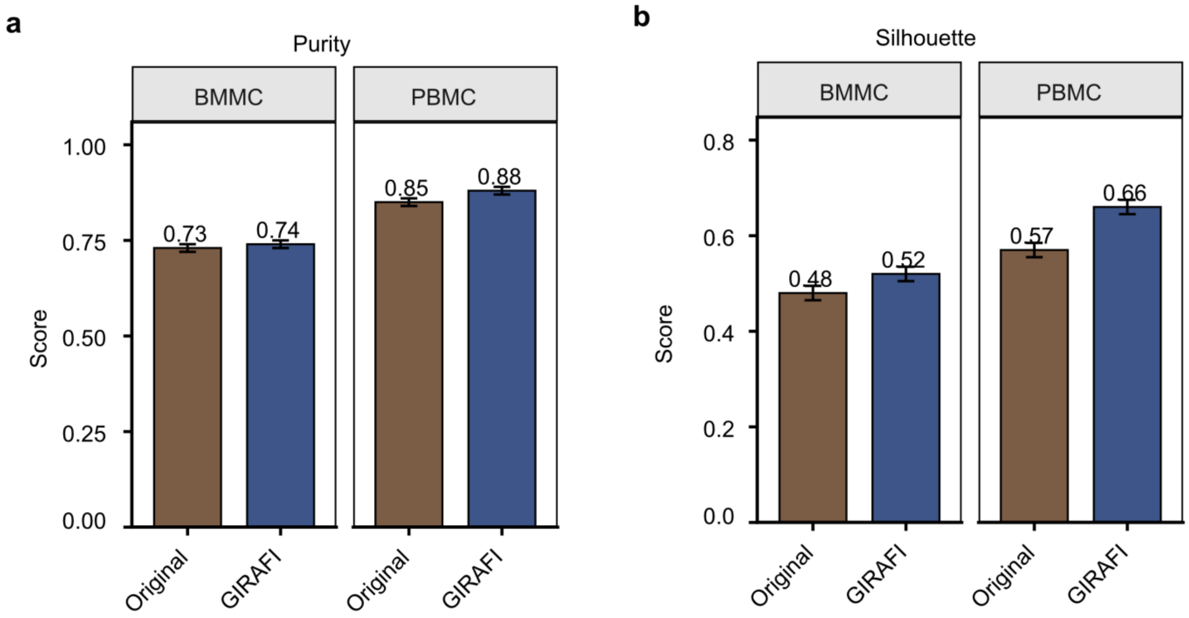

**Supplementary Fig. 10: Cluster quality metrics in BMMC and PBMC after imputation.**

Bar plots summarize two complementary structure metrics computed within BMMC and PBMC for the Original and GIRAFI matrices.

**a,** Cluster purity after unsupervised clustering. For each dataset and method, cells were clustered without using the reference labels, and the resulting cluster assignments were compared with the annotated cell-type labels. Purity was calculated as the fraction of cells belonging to the dominant annotated cell type within each inferred cluster, averaged across clusters; higher values indicate that inferred clusters are more consistently composed of a single annotated cell type.

**b,** Mean silhouette score, reflecting within-group cohesion relative to between-group separation in the embedding/feature space used for evaluation.

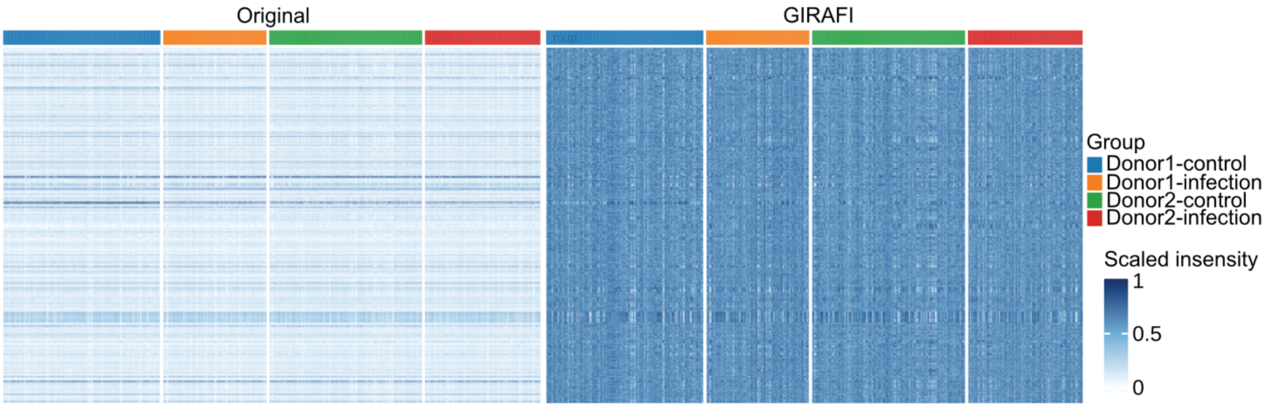

**Supplementary Fig. 11: GIRAFI increases within-group coherence in the astrocyte dataset across donors and conditions.**

Protein-by-cell heat maps comparing the Original and GIRAFI matrices, with columns grouped by donor condition subsets.

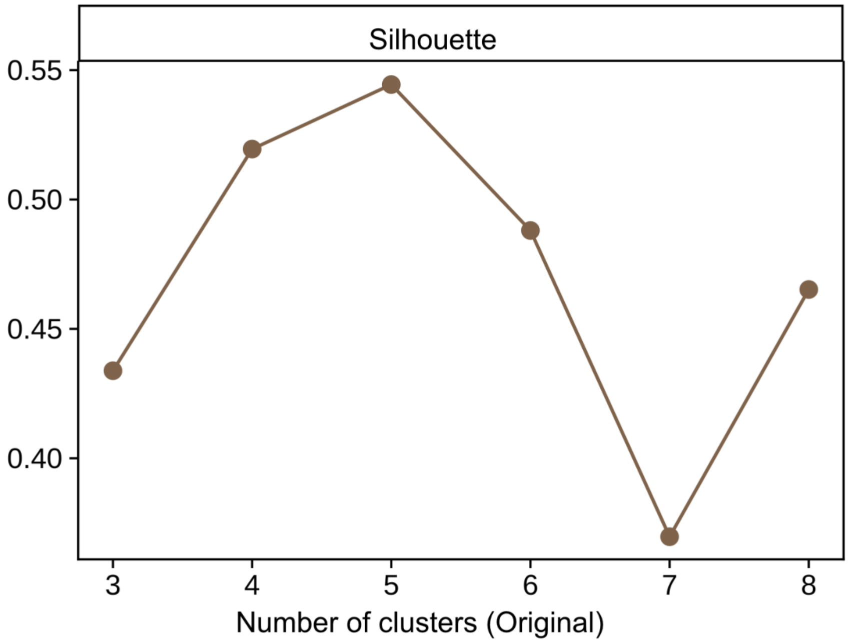

**Supplementary Fig. 12: Silhouette analysis across candidate cluster numbers in the Original astrocyte representation.**

Silhouette coefficients computed across candidate cluster numbers for the Original representation.

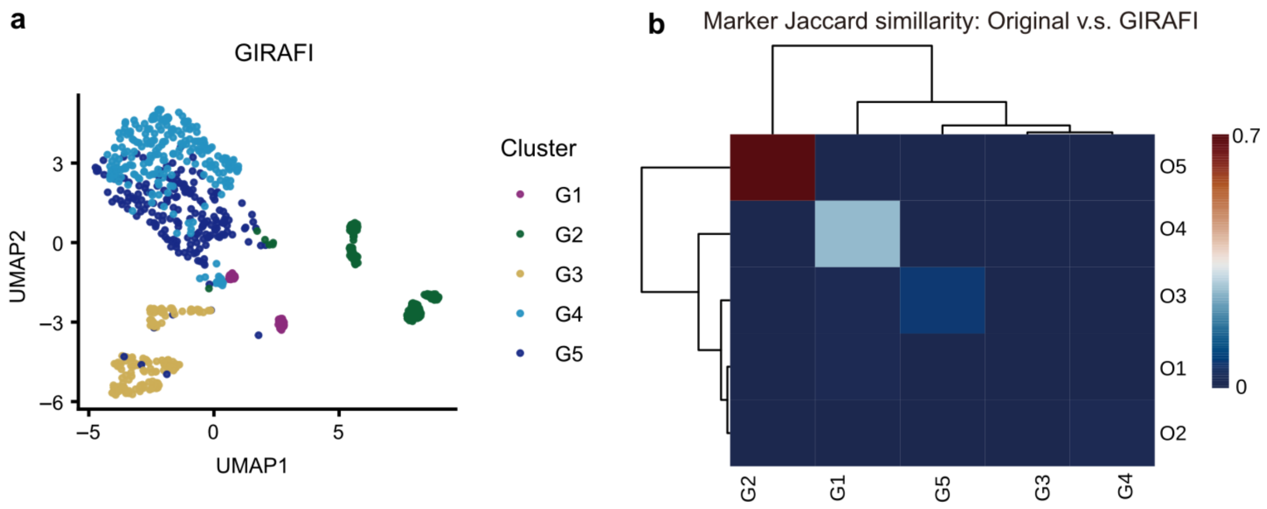

**Supplementary Fig. 13: Matched-granularity clustering and cross-method correspondence between the Original and GIRAFI astrocyte representations.**

**a, UMAP embedding of the GIRAFI matrix annotated with fixed cluster assignments.** Cells are coloured by cluster identity, with five GIRAFI-derived clusters denoted G1-G5.

**b, Cross-method cluster correspondence between the Original and GIRAFI matrices.** The heat map shows Jaccard similarity between marker protein sets, with rows representing Original clusters (O1-O5) and columns representing GIRAFI clusters (G1-G5). Rows and columns are ordered by hierarchical clustering, and warmer colours indicate greater marker-set overlap.

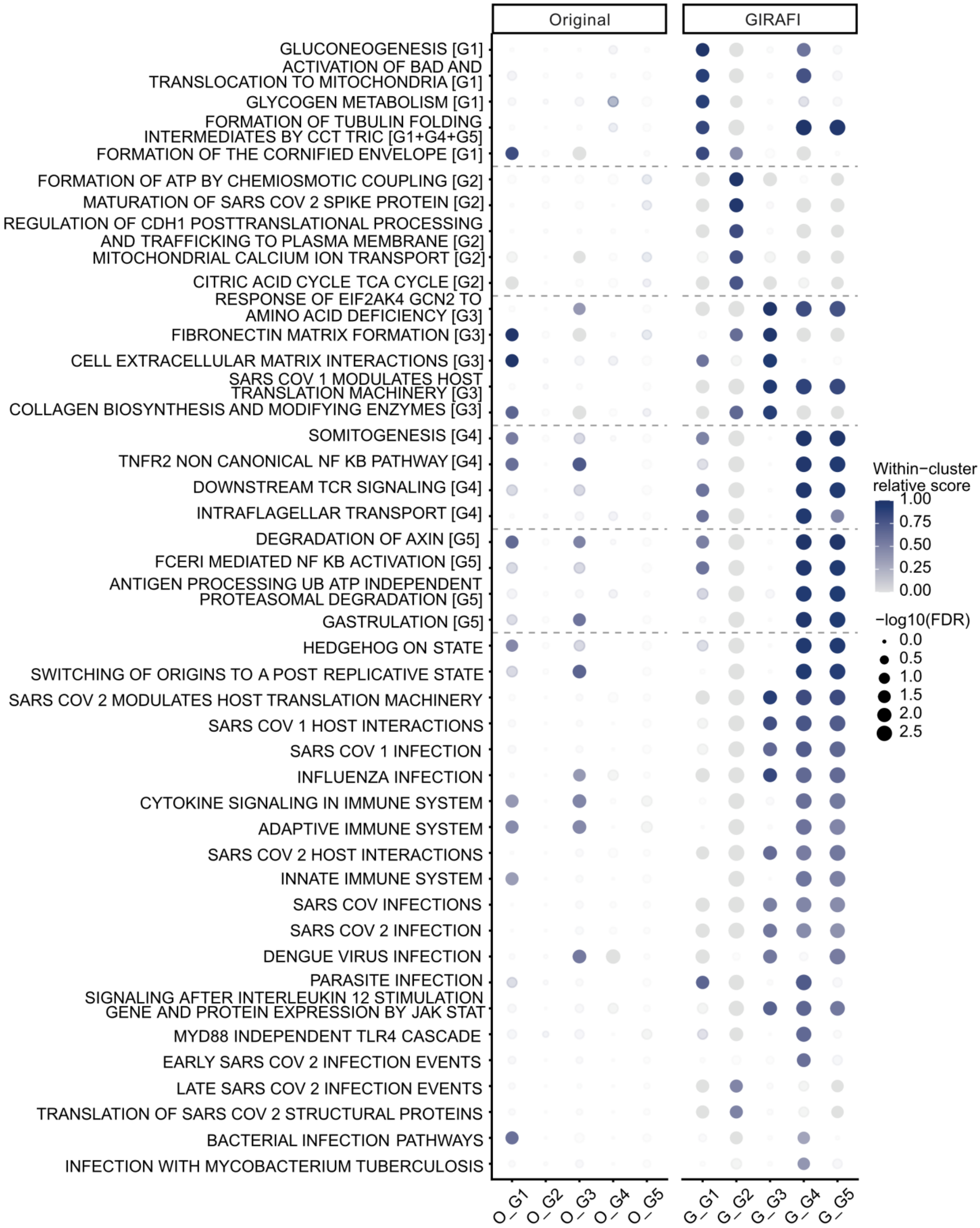

**Supplementary Fig. 14: Reactome selected contrast terms across fixed-k clusters (FGSEA).**

Dot plot shows per-cluster Reactome enrichment profiles for the Original and GIRAFI matrices using fixed-k clusters (O1-O5; G1-G5). Reactome terms were restricted to top enriched terms in clusters, as well as infection/viral annotations and further de-redundant, then selected as GIRAFI-enriched (high in GIRAFI, low/absent in Original) based on the between-method delta score from the all-clusters FGSEA summary. Dot size encodes -log10(FDR) and dot color encodes the normalized enrichment score (NES; positive values indicate enrichment toward the top of the ranked list, negative values toward the bottom).

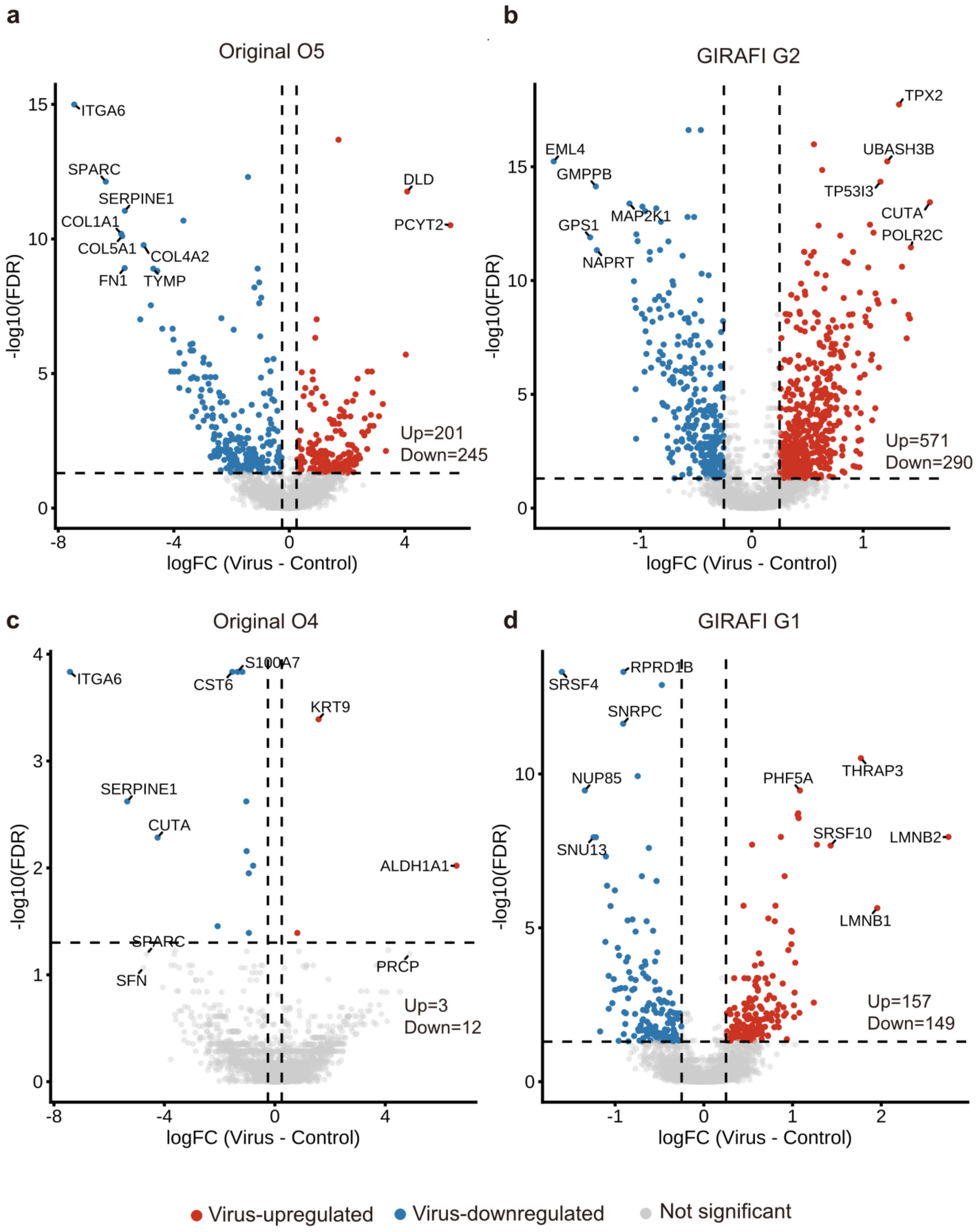

**Supplementary Fig. 15: Matched-state volcano plots comparing virus-versus-control signals in the Original and GIRAFI representations.**

Example "shared-state" cluster pairs (marker Jaccard values are reported in the titles) illustrating within-cluster differential expression between Virus and Control conditions in the Original and GIRAFI data. Volcano plots compare virus-infected versus control cells within each matched cluster pair; dashed lines indicate the significance and effect-size thresholds. Points are colored by direction of change (virus-up, control-up, or not significant), selected proteins are labeled, and the numbers of significantly up- and down-regulated proteins are annotated in each panel.

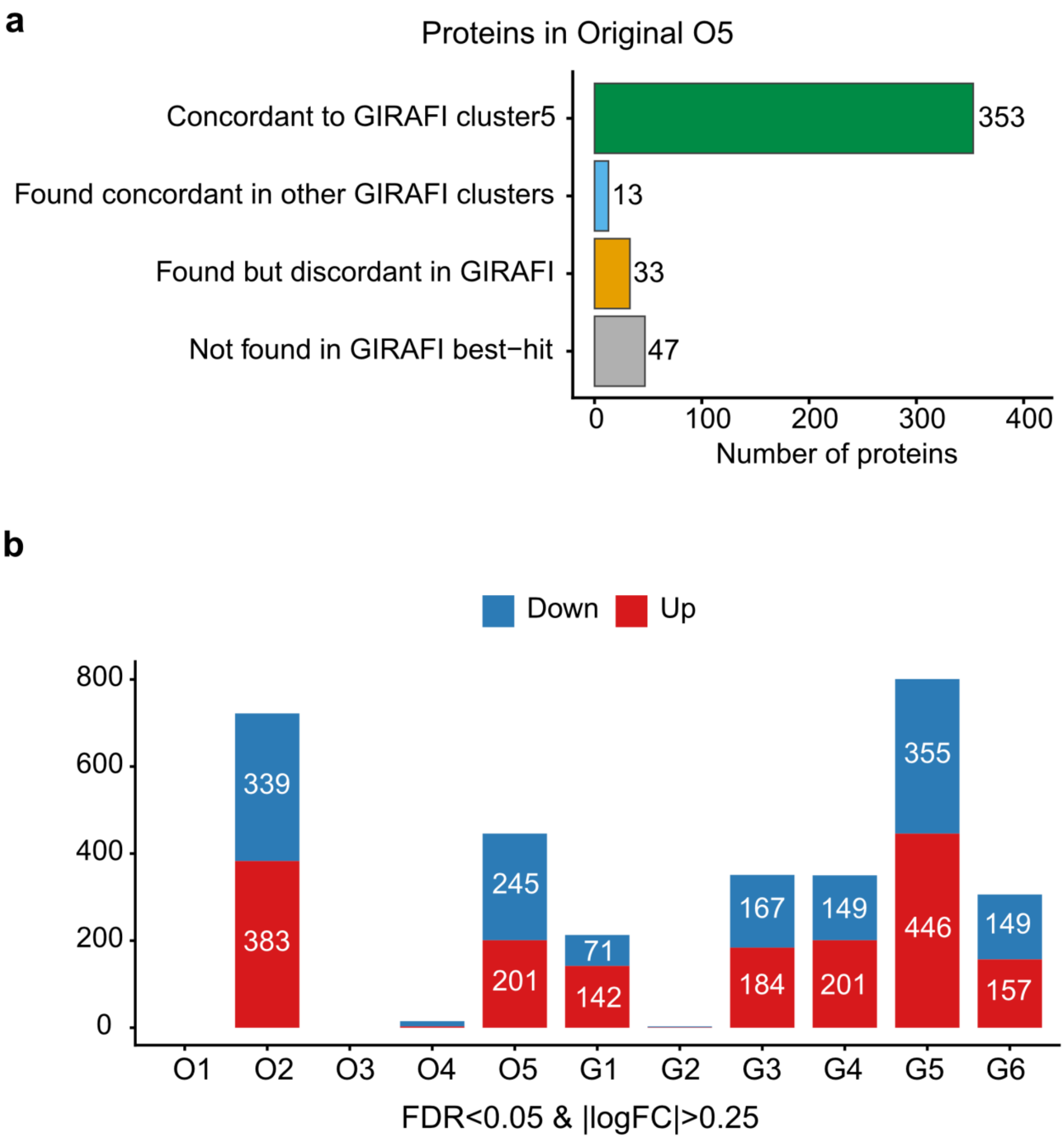

**Supplementary Fig. 16: Differentially expressed protein redistribution and direction consistency between Original and GIRAFI cluster-level signals**

**a**, Direction concordance for a representative comparison: proteins that are significant in the Original analysis (strict threshold shown in the panel title) are stratified into four categories based on whether they (i) show the same direction in the matched GIRAFI cluster, (ii) reappear with the same direction but in a different GIRAFI cluster, (iii) are found but with discordant direction, or (iv) are not recovered among the GIRAFI best-hit signals.

**b**, The number of significant differentially expressed proteins (Virus vs Control) is shown for each Original cluster (O1-O5) and each GIRAFI cluster (G1-G6), split into Up and Down counts (FDR and |log FC| thresholds as annotated).

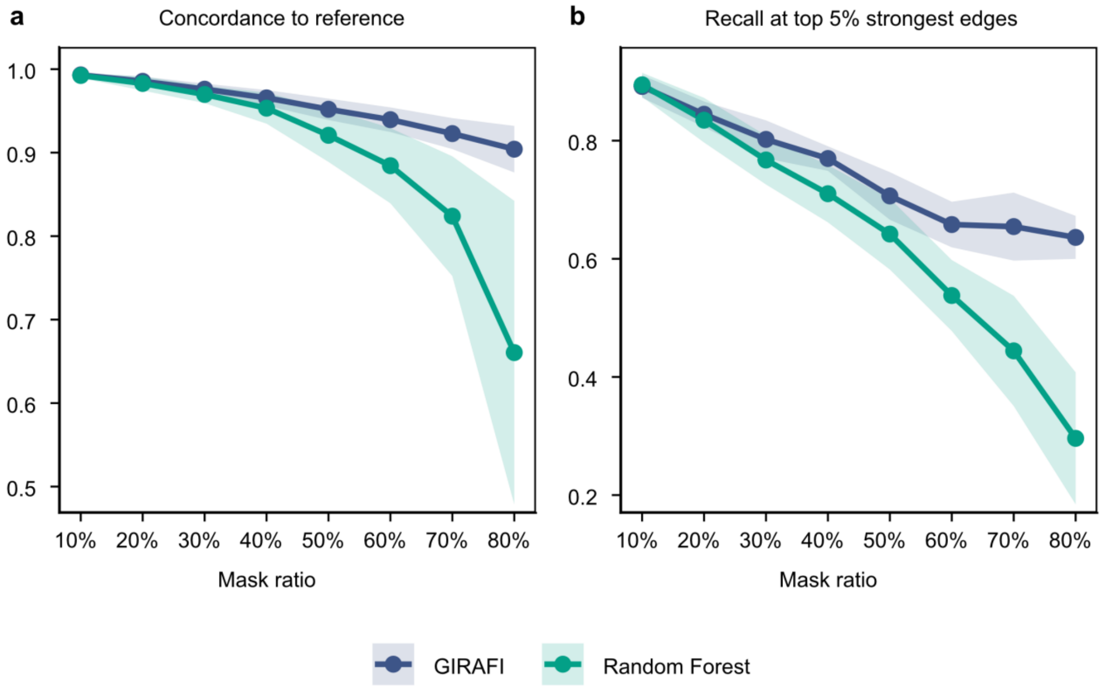

**Supplementary Fig. 17: GIRAFI preserves PPI edge-level correlation structure under increasing masking.**

PPI edge-level recovery was evaluated in a virally infected astrocyte dataset (two donors × two conditions) by a mask-and-recover design. Starting from the complete matrix, 10-80% of observed entries were randomly masked and then imputed with either Random Forest or GIRAFI. For each mask ratio, we computed an edge-correlation profile by estimating Spearman correlations across cells for curated PPI-linked protein pairs. Top, global concordance to the reference profile, measured as the correlation between the imputed and reference edge-correlation vectors. Bottom, recovery of the strongest interaction-consistent signals quantified by Recall at top5%, defined as the fraction of reference top-5% edges (ranked by absolute correlation) that are also present among the imputed top-5% edges.

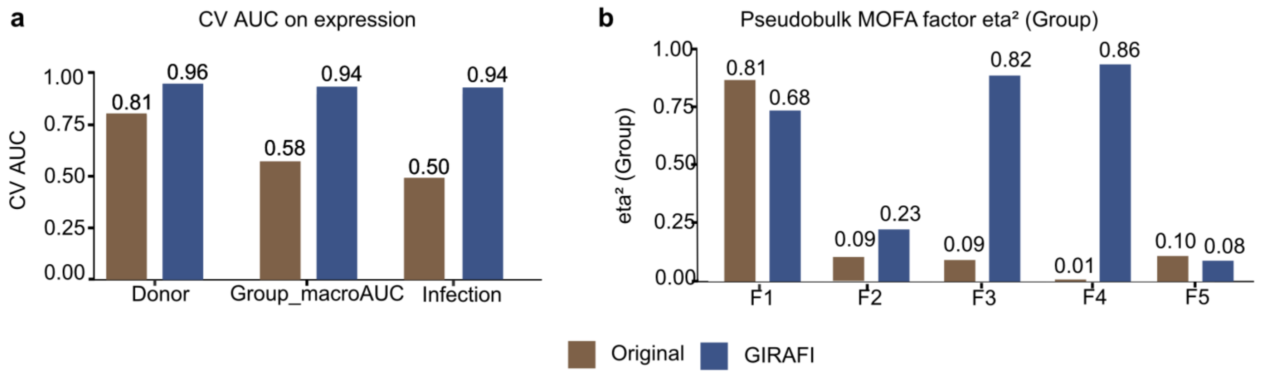

**Supplementary Fig. 18: Predictability of metadata and group-associated factor structure under GIRAFI versus Original**

**a,** Cross-validated AUC for predicting metadata targets from expression profiles. Cross-validated AUC (expression residualized as indicated) for predicting metadata targets, comparing Original and GIRAFI: donor identity, group (macro-AUC), and infection status.

**b,** Variance explained by group, measured as $\eta^2$, for pseudobulk MOFA factors. Variance explained ($eta^2$) by group for pseudobulk MOFA factors (F1-F5) computed under each method (Original vs GIRAFI), summarizing how strongly group structure is captured at the factor level.

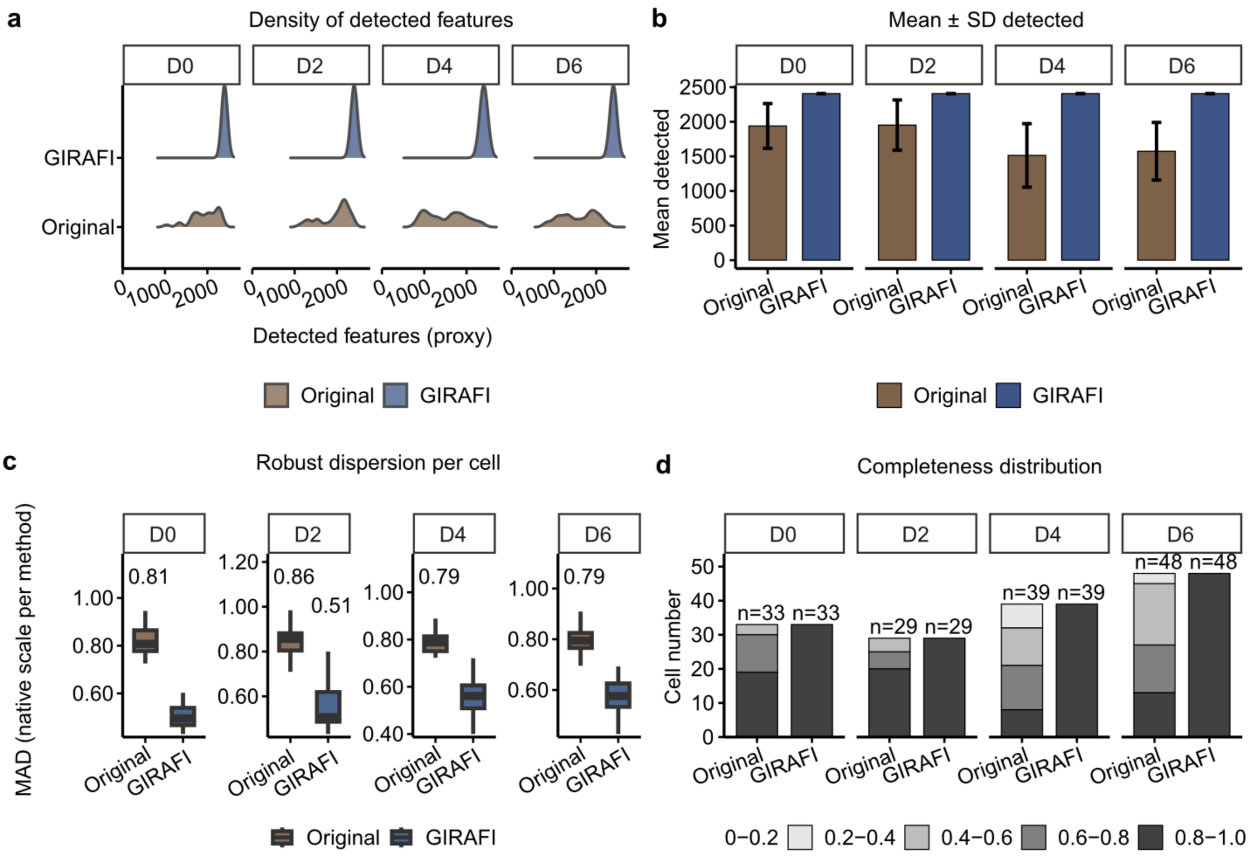

**Supplementary Fig. 19: Quality-control summary for the PC12 time-course: Original versus GIRAFI.**

**a, Distributions of detected protein features per cell across the PC12 time course.** Detection-proxy distributions, defined as the number of detected protein features per cell, are shown for each timepoint (D0, D2, D4, and D6) in the Original and GIRAFI matrices.

**b, Mean ± SD of detected protein features per cell across timepoints.** Bars summarize the mean number of detected protein features per cell for each timepoint in the Original and GIRAFI matrices, and error bars indicate the standard deviation.

**c, Robust per-cell dispersion across timepoints.** Per-cell dispersion was quantified by the median absolute deviation (MAD) after outlier removal. For the Original matrix, dispersion was computed from log_2(raw intensity) values without a pseudocount. For the GIRAFI matrix, dispersion was computed from the model output on its native scale without range projection.

**d, Completeness distributions across cells over the PC12 time course.** Stacked bars show the number of cells falling into bins of observed-protein fraction for each timepoint and method. Numbers above the bars indicate the total number of cells in each group.

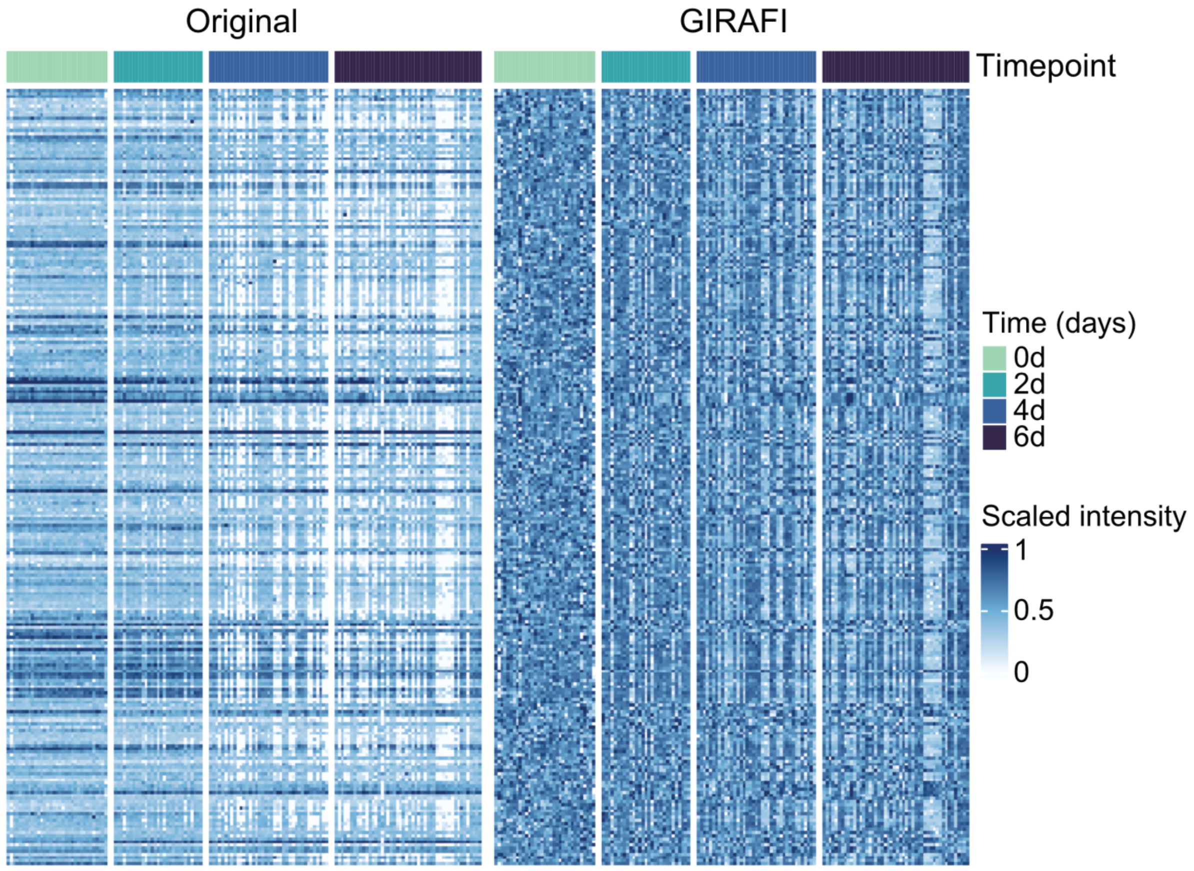

**Supplementary Fig. 20: Global expression structure across timepoints in Original and GIRAFI matrices.**

Heatmaps show protein-by-cell expression for the Original matrix (left) and GIRAFI (right), with cells grouped by timepoint and values scaled to 0-1 for visualization (Original: zeros held fixed; GIRAFI: scaled 0-1).

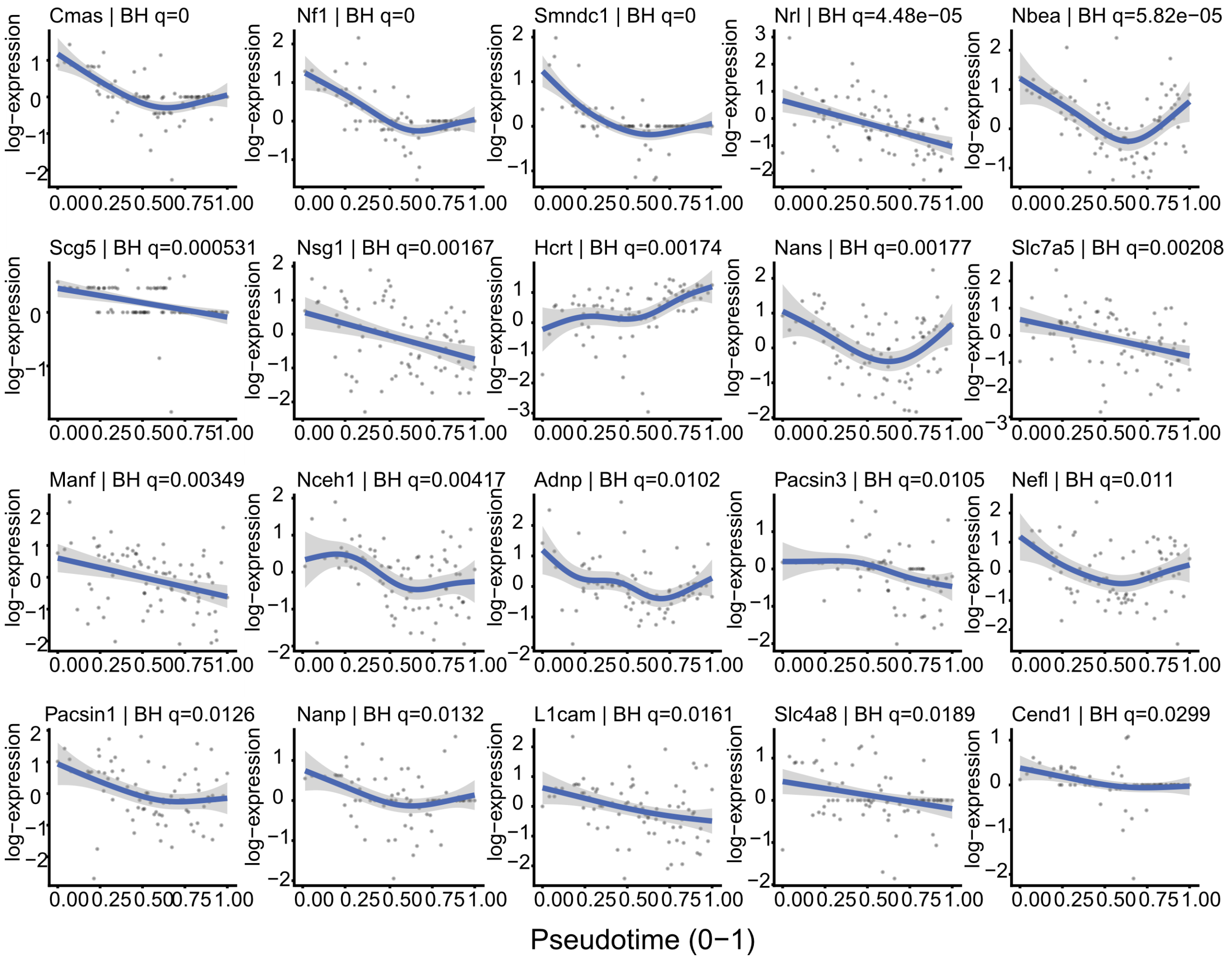

**Supplementary Fig. 21: Representative proteins show graded expression changes along pseudotime.**

Each small-multiple panel shows one protein's expression as a function of pseudotime (x-axis; scaled to 0-1). Grey points represent single cells (or single-cell profiles) and the blue curve depicts a smoothed trend fitted across pseudotime; the light-blue ribbon indicates the uncertainty around the fitted trend (confidence band). The y-axis shows log-transformed expressions. Panel titles report the protein identifier and the Benjamini-Hochberg multiple-testing-adjusted q-value (BH q) for association with pseudotime, highlighting proteins with monotonic as well as transient/inflection-point patterns. (q = 0 indicates an adjusted value below numerical display precision.)

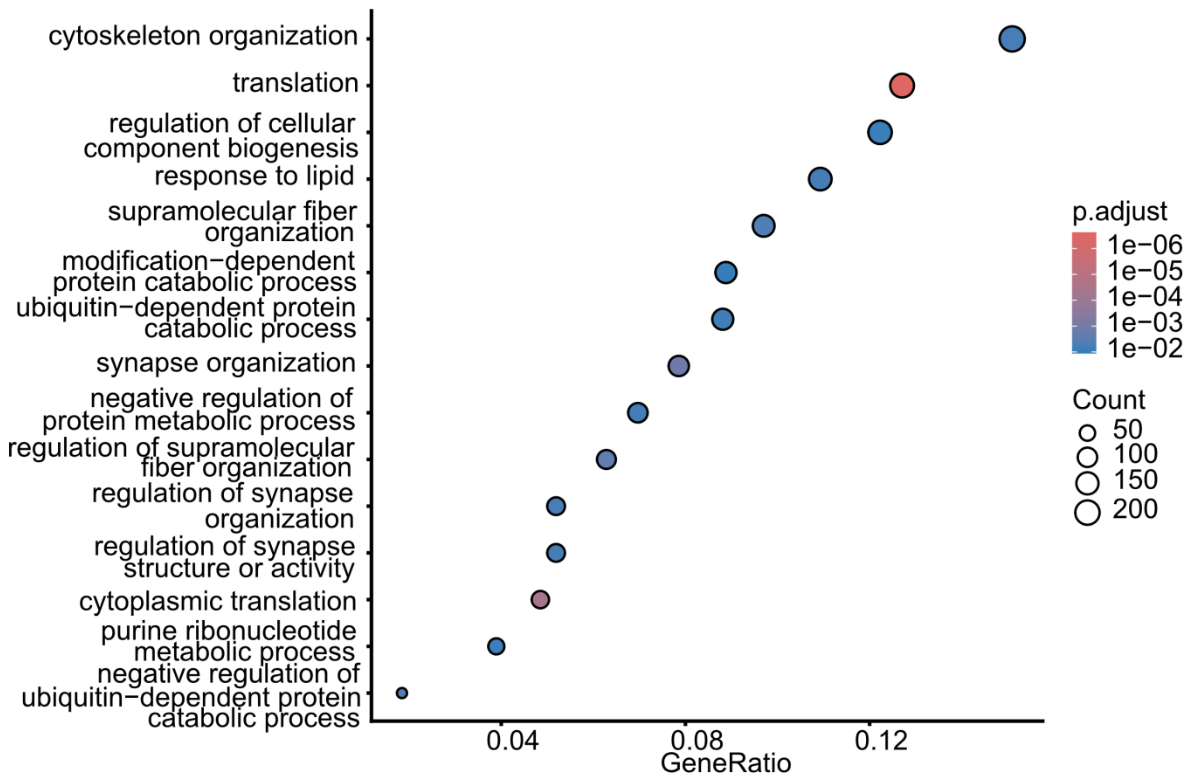

**Supplementary Fig. 22: Functional interpretation of pseudotime-associated proteins.**

Over-representation analysis (ORA) of pseudotime-increasing proteins using GO Biological Process terms; point size indicates gene count and colour represents adjusted significance.

**Supplementary Note**

**Data preprocessing**

SCP DIA-NN output was processed using a unified pipeline that supports both older formats and newer exports with protein annotations. Contaminant entries were removed based on a standard contaminant list, and rows without valid gene names were excluded.

Cells with no detected proteins and proteins with no observed values were removed before downstream analysis. Cells were then filtered using two quality-control metrics: total raw intensity and the number of detected proteins per cell. Total raw intensity was calculated as the sum of all observed protein intensities in each cell, and cells with unusually low or high total intensity were removed using either quantile-based or absolute thresholds, depending on the dataset. The detected-protein count was calculated as the number of proteins observed in each cell. Unless otherwise specified, cells were retained only if they contained more than 500 and no more than 4,000 detected proteins. These cutoffs were adapted to each experimental context, including acquisition depth and dataset-specific sparsity, to remove low-quality outliers while preserving the expected cell-type structure and sufficient cell numbers for downstream analyses.

Proteins were then filtered by detection frequency. Unless otherwise specified, proteins were retained if they were observed in at least 20% of cells. Remaining intensity values were log-transformed after adding a small pseudocount, defined as half of the minimum positive observed intensity. Log-transformed intensities were then standardized separately for each protein by subtracting the protein-wise mean and dividing by the protein-wise standard deviation. Missing entries were retained as missing values rather than being replaced during this preprocessing step. After transformation and standardization, proteins with more than 85% missing values were removed unless otherwise specified.

**Adjusted Rand Index (ARI)**

We quantified agreement between predicted clusters and known labels using the adjusted Rand index (ARI), which measures pairwise consistency of assignments while correcting for chance. Formally, for a contingency table $n_{ij}$ between clusters $i$ and labels $j$, ARI is defined as

$$\mathrm{ARI} = \frac{\sum_{ij}\binom{n_{ij}}{2} - \left[\sum_i\binom{a_i}{2}\sum_j\binom{b_j}{2}\right]/\binom{n}{2}}{\frac{1}{2}\left[\sum_i\binom{a_i}{2} + \sum_j\binom{b_j}{2}\right] - \left[\sum_i\binom{a_i}{2}\sum_j\binom{b_j}{2}\right]/\binom{n}{2}}.$$

where $a_i = \sum_j n_{ij}$, $b_j = \sum_i n_{ij}$, and $n$ is the total number of cells. ARI ranges from 0 (chance-level agreement) to 1 (perfect agreement), and may be negative when agreement is worse than expected by chance.

### Silhouette score

We assessed cluster separation using the silhouette coefficient, which compares within-cluster cohesion to nearest-cluster separation. For each cell $i$, let $a(i)$ be the average distance from $i$ to all other cells in its assigned cluster, and let $b(i)$ be the minimum (over all other clusters) of the average distance from $i$ to cells in that cluster. The silhouette for $i$ is

$$s(i) = \frac{b(i) - a(i)}{\max\{a(i), b(i)\}}.$$

The overall silhouette score is the mean of $s(i)$ over all cells. Values close to 1 indicate well-separated clusters, values near 0 indicate overlap, and negative values indicate that cells are, on average, closer to a different cluster than to their assigned cluster.

### Precision, recall, and F1-score

We evaluated label recovery using precision, recall, and F1-score, computed from the counts of true positives (TP), false positives (FP), false negatives (FN), and true negatives (TN).

### Binary classification

Precision quantifies the fraction of predicted positives that are correct:

$$\text{Precision} = \frac{\text{TP}}{\text{TP} + \text{FP}}.$$

Recall (sensitivity) quantifies the fraction of true positives that are recovered:

$$\text{Recall} = \frac{\text{TP}}{\text{TP} + \text{FN}}.$$

The F1-score is the harmonic mean of precision and recall:

$$\text{F1} = \frac{2\ \text{Precision}\ \text{Recall}}{\text{Precision} + \text{Recall}} = \frac{2\ \text{TP}}{2\ \text{TP} + \text{FP} + \text{FN}}.$$

### Multi-class classification

For multi-class settings, we computed class-wise precision/recall/F1 under a one-vs-rest scheme and reported aggregated scores.

- **Macro-average** (treats all classes equally):

$$
\begin{aligned}
\text{Precision}_{\text{macro}} &= \frac{1}{\text{K}}\sum_{k=1}^{\text{K}} \text{Precision}_{\text{k}}\,, \\
\text{Recall}_{\text{macro}} &= \frac{1}{\text{K}}\sum_{k=1}^{\text{K}} \text{Recall}_{\text{k}}\,, \\
\text{F1}_{\text{macro}} &= \frac{1}{\text{K}}\sum_{k=1}^{\text{K}} \text{F1}_{\text{k}}\,.
\end{aligned}
$$

- **Weighted macro-average** (weights by class support $n_k$):

$$
\text{F1}_{\text{weighted}} = \sum_{k=1}^{K} \frac{n_k}{n}\, \text{F1}_k \text{ (analogously for precision/recall).}
$$

- **Micro-average** (pools decisions across classes; common for imbalanced data):

$$
\begin{aligned}
\text{Precision}_{\text{micro}} &= \frac{\sum_{k=1}^{\text{K}} \text{TP}_{\text{k}}}{\sum_{k=1}^{\text{K}} (\text{TP}_{\text{k}} + \text{FP}_{\text{k}})}, \\
\text{Recall}_{\text{micro}} &= \frac{\sum_{k=1}^{\text{K}} \text{TP}_{\text{k}}}{\sum_{k=1}^{\text{K}} (\text{TP}_{\text{k}} + \text{FN}_{\text{k}})}, \\
\text{F1}_{\text{micro}} &= \frac{2\,\text{Precision}_{\text{micro}}\text{Recall}_{\text{micro}}}{\text{Precision}_{\text{micro}} + \text{Recall}_{\text{micro}}}.
\end{aligned}
$$
